# Recognition and remodelling of nucleosomes and hexasomes by the human INO80 complex

**DOI:** 10.1101/2025.08.25.671740

**Authors:** Priyanka Aggarwal, Manmohan Sharma, Stephan Woike, Franziska Kunert, Manuela Moldt, Karl-Peter Hopfner

## Abstract

The ATP-dependent INO80 chromatin remodeller slides and repositions nucleosomes to shape and maintain chromatin around gene regulatory elements and replication origins. Recent work uncovered capabilities of yeast and fungal INO80 to bind and slide hexasomes, but whether this is a universal feature is unknown. Here, we show that also human INO80 slides hexasomes as efficiently as H2A and H2A.Z nucleosomes. By determining a variety of structures of human INO80 bound to canonical and H2A.Z nucleosomes as well as hexasomes, we reveal a predominantly topological sensing of nucleosomal species with at least three positions depending on entry DNA unwrapping. INO80 spin-rotates around the wheel like nucleosomal core particle, with the position determined by the Snf2 ATPase that binds the tangentially protruding entry DNA at various degrees of unwrapping. Acidic patch binding by IES2 can differentiate between different nucleosomal species, is important for nucleosome but not hexasome sliding, and can sense unwrapped exit DNA. These findings provide structural and mechanistic insights into how human INO80 remodels diverse chromatin substrates in a topology driven manner.

## Introduction

Chromosomal DNA resides in the nucleus in the form of chromatin, a dynamic and variable complex of DNA and associated proteins, primarily histones. The major building blocks of chromatin are nucleosomes, which consist of ∼146 base pairs (bp) of DNA wrapped around a histone octamer in ∼1.7 left-handed superhelical turns (1). The histone octamer consists of two copies each of the canonical histones H2A, H2B, H3, and H4, or their respective variants and isoforms (2,3). Nucleosomes package and protect the DNA, however they are also important carriers of epigenetic information, which is encoded in their local chromosomal position, histone composition and chemical modification (4). The position and epigenetic states of nucleosomes help govern transcription and gene regulation, mark chromosomal loci and help conduct other DNA associated processes such as replication, recombination and repair (5–7).

The nucleosomal landscape along DNA is shaped by the collective action of ATP-dependent chromatin remodellers along with numerous other factors (5–7). Chromatin remodellers use the energy of ATP hydrolysis to slide, position or edit (i.e., exchange histone variants) nucleosomes, or evict histones altogether. They are grouped into, but are not limited to, four main families, denoted INO80, SWI/SNF, ISWI and CHD. Chromatin remodellers display diverse domain and subunit architectures, but share an ATP dependent motor domain, which belongs to the Snf2 family among superfamily 2 helicases/translocases. Using cycles of ATP binding and hydrolysis, Snf2 ATPases translocate DNA and/or alter local DNA shape, which serves as the underlying chemo-mechanical activity governing the diverse remodelling reactions (5–7).

The INO80 family consists of INO80 and SWR1 type complexes (8,9). Both remodellers are involved in shaping chromatin, particularly around nucleosome-depleted or nucleosome-free regions (NDRs/NFRs), such as promoters and origins of replication. They can sense extended stretches of extranucleosomal entry DNA. Despite this shared ability, they catalyze distinct nucleosome reconfigurations: SWR1 primarily edits nucleosomes by exchanging histone variants, whereas INO80 mainly repositions nucleosomes along the DNA (9,10). INO80 can slide nucleosomes and place them at the boundaries of NDRs *in vitro*. INO80 can also generate nucleosomal arrays outwards from these NDRs (11,12). SWR1, on the other hand, incorporates the H2A variant H2A.Z to replace canonical H2A in nucleosomes (13–15). In humans, the same function is carried out by SRCAP and TIP60 complexes (16). H2A.Z is enriched at promoters, enhancers and origins of replication and has pleiotropic, context and species dependent functions including roles in transcription regulation, DNA repair, and others (17–20). Histone exchange activity has also been reported for yeast INO80 *in vitro*, with a preference to exchange H2A.Z to H2A, but its physiological role is not yet fully understood (21,22).

Yeast and fungal INO80 can also slide subnucleosomal particles, such as hexasomes (23–25). Hexasomes are characterized by loss of one histone dimer, typically H2A/H2B (25). They are proposed to arise when RNA polymerase II transcribes through the nucleosome. Depending on the orientation they either block or allow transcription by RNAPII *in vitro* (26). Although our understanding of hexasomes’ roles or their presence in various genomic processes *in vivo* remains limited, their potential occurrence as transient intermediates or products in numerous nucleosomal activities or remodelling reactions underscores the importance of understanding how remodellers interact with the subnucleosomal particles and non-standard nucleosome variants.

Previous biochemical studies have revealed that the INO80 complex possesses a modular structure with more than 15 distinct subunits (27). The largest polypeptide, INO80, harbors the Snf2 motor domain and also functions as scaffold on which the other modules or subunits assemble (28,29). The N-terminal “N-module” (human subunits NFRKB, TFPT/Amida, MCRS1, UCHL5, INO80D, INO80E assembled at the N-terminal region of INO80) is very divergent in evolution and its function is not well understood, since it is dispensable for *in vitro* remodelling activity (28,30). The central “A-module” contains nuclear actin along with actin related proteins ACTB, ACTL6A and ACTR8 and the zinc finger containing transcriptional repressor YY1(31). The A-module binds extranucleosomal entry DNA and helps couple the motor activity to nucleosome sliding. INO80’s C-terminal region forms the nucleosome core particle (NCP) mobilizing module and harbours the Snf2 domain (Ino80^motor^), along with nucleosome binding subunits ARP5 (actin-related protein 5), /IES6 subunit (encoded by ACTR5/ INO80C), IES2 (Ino eighty subunit 2) and the assembly chaperone heterohexamer AAA^+^ ATPases RuvBL1/RuvBL2 (28,29,31).

While structures of members from all major families have been visualized bound to canonical nucleosomes (14,16,29,32,33), considerably less is known how remodellers interact with non-canonical nucleosomes containing histone variants or subnucleosomal particles. Recently, yeast and fungal INO80 complexes and SWRI bound to hexasomes have been visualised by cryogenic electron microscopy (cryo-EM) (23,24,34). Yeast and fungal INO80 complexes bind hexasomes in a spin-rotated orientation compared to nucleosome binding, with the H2A–H2B dimer on the DNA-entry side absent. Lack of acidic patch (negatively charged surface region on the nucleosome, located mainly on histones H2A and H2B) interaction due to the missing H2A/H2B dimer and ∼35 base pairs of unwrapped entry-side DNA led to a different binding mode through formation of new interfaces between yeast and fungal INO80 and the hexasome in comparison with nucleosome interactions (23,24). However, this mode of subnucleosomal particle binding does not appear to be a general feature of chromatin remodellers. The Chd1 remodeller requires an H2A/H2B at the DNA entry side of the nucleosome, and slides hexasomes in the opposite way than observed for INO80 *in vitro* (35). Furthermore, since some nucleosome recognition features, in particular the ARP5/ACTR5 insertion element that binds the exposed histone surface of the hexasome in yeast and fungal INO80 is missing in human INO80, it is yet unclear to what extent sliding of non-canonical nucleosomes is even present in the mammalian system.

Here, we show that the human delta-N INO80 complex (C– and A-module, lacking the N-terminal module) can slide hexasomes as efficiently as nucleosomes, similar to its yeast and fungal orthologs. This demonstrates an evolutionary conserved apparent indifference to nucleosomal and hexasomal species with respect to basal sliding activity. The structures of the human INO80 complex bound to hexasomes, H2A.Z-nucleosomes and canonical nucleosomes in different states shows a high capability for spin-rotation, whereby essentially the location of entry DNA and its interaction with the Ino80^motor^ domain determine placement of the remodeller.

## Materials and methods

### Expression and purification of delta-N INO80

The open reading frames (ORFs) of subunits of *H. sapiens* INO80 (C-module comprising RuvBL1, RuvBL2, ARP5, IES2, IES6, and Ino80^270-1552^ having a 6X Histidine tag at the N-terminus and 2X Streptavidin tag at the C-terminus) and A-module (ARP8, Actin, ARP4 and YY1) were optimized for insect cell expression and ordered from GeneArt (Thermo Fisher Scientific). The gene cassettes for C and A-modules were assembled separately on two pBIG2ab vectors employing the biGBac cloning system (36). The baculoviruses generation was done in Sf21 insect cells (*S. frugiperda,* Thermo Fisher Scientific, #11497013), after which the complexes were recombinantly expressed in High Five insect cells (*Trichoplusia ni*; Invitrogen, #B85502) by adding the two viruses at a ratio of 1:100 (volume virus: medium) to three liters of insect cell culture. Cells were cultured for 60 hours at 27 °C. Cells were harvested by centrifugation at 4 °C.

The complex was purified according to the previously published protocol (37). Briefly, the complex was purified by Ni-NTA affinity followed by Strep-Tag affinity and HiTrap Q (1 ml) chromatography. Initially, the cell pellet was resuspended in lysis buffer (50 mM Tris, pH 8.0, 500 mM NaCl, 1 mM TCEP, 2 mM benzamidine-HCl, 10% (v/v) glycerol) supplemented with 10 µl BaseMuncher nuclease (Abcam, ab270049) and one EDTA-free protease inhibitor tablet (Sigma-Aldrich). The lysate was gently sonicated for 2 minutes (duty cycle, 50%; output control, 5). After sonication, the lysate was clarified by centrifugation at 30,500 g for 1 hour at 4 °C, filtered through a 0.22 µm Millipore filter. The lysate was loaded onto a pre-equilibrated 5 ml HisTrap FF column (Cytiva) using a peristaltic pump. The column was washed with 10 column volumes of His-buffer A (50 mM Tris, pH 8.0, 250 mM NaCl, 1 mM TCEP, 10% (v/v) glycerol), and the complex was eluted with five column volumes of His-buffer B (His-buffer A with 250 mM imidazole). The eluted complex was immediately loaded onto pre-equilibrated Strep-Tactin®XT Superflow® resin (IBA Lifesciences, 3 ml slurry) and incubated for 25 minutes with gentle rolling at 4 °C.

The resin was then washed with Strep-buffer A (50 mM Tris, pH 8.0, 200 mM NaCl, 1 mM TCEP, 10% (v/v) glycerol). After washing, the complex was eluted iteratively in Strep-elution buffer (Strep-buffer A supplemented with 5 mM desthiobiotin, Sigma-Aldrich) over a period of 60 minutes. The eluate was subsequently loaded onto a HiTrap Heparin HP column (Cytiva), and the protein was eluted using a linear salt gradient from 100% HiTrap-Q buffer A (50 mM Tris, pH 8.0, 1 mM TCEP, 10% (v/v) glycerol) to 100% HiTrap-Q buffer B (50 mM Tris, pH 8.0, 1 M NaCl, 1 mM TCEP, 10% (v/v) glycerol) over 12 column volumes. The fractions were analyzed by SDS-PAGE and those with the highest purity were pooled together.

The protein complex was concentrated to 5 mg/ml using centrifugal filters (Centricon, 50 kDa cutoff, Millipore) and either used for vitrification on the same day or dialyzed in storage buffer (50 mM Tris, pH 8.0, 250 mM NaCl, 10% (v/v) glycerol, 1 mM TCEP) before being flash-frozen in liquid nitrogen.

INO80 mutants were generated by site-directed mutagenesis PCR and were expressed and purified using the same protocol as the wild-type protein. List of primers used for preparation of INO80 mutants are listed in **Table 1**.

**Table 1.**
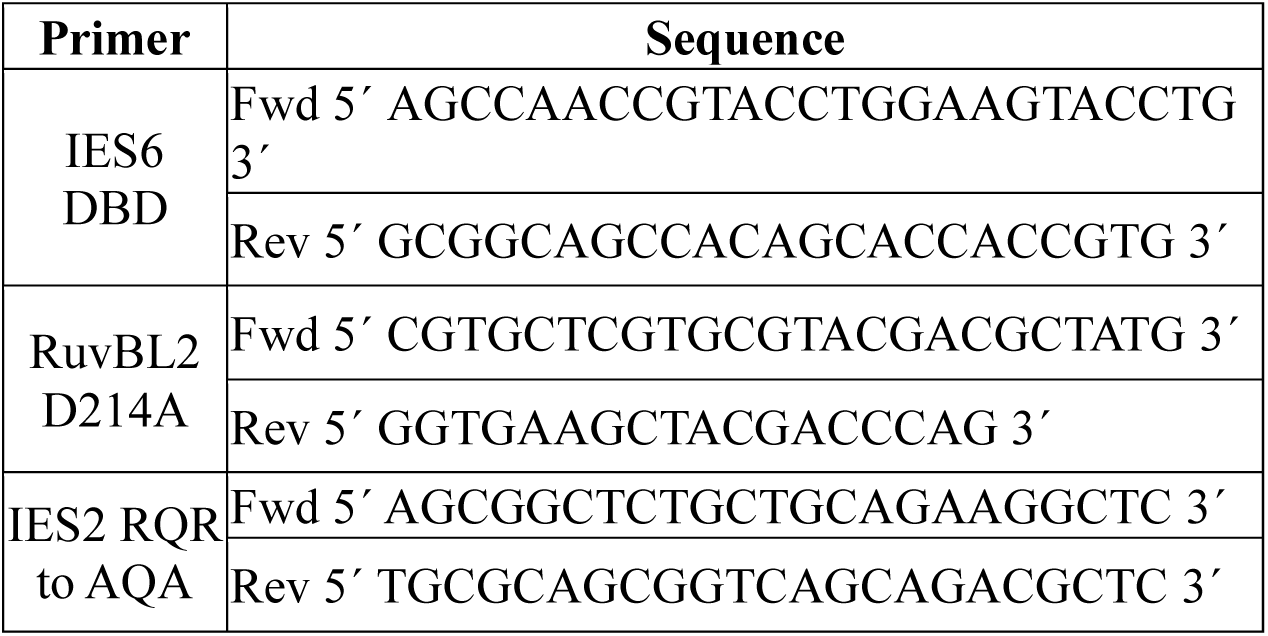
– DNA primers used for generating the mutants.

### Preparation of nucleosomes and hexasomes

Canonical human histones were obtained from the histone source at Colorado State University, USA, and unfolded in a buffer containing 7 M guanidinium chloride, 20 mM Tris (pH 7.5), and 1 mM DTT at room temperature for 30 minutes with continuous stirring. Following this, histone dimers, tetramers, and octamers were refolded according to methods described in previous publications (38). The histone H2A mutant (E61A/E64A/D72A/D90A) was also expressed and purified as previously reported (29)(38). Widom 601-DNA (39) with varying linker lengths was amplified via polymerase chain reaction (PCR) and purified using anion exchange chromatography. The purified DNA was then concentrated under vacuum and stored at –20°C. The details of the primers used are provided in **Table 2**. Nucleosomes were reconstituted by mixing the octamer and DNA in a 1:1 ratio at 2 M NaCl, with the salt concentration gradually reduced to 50 mM over 16 hours at 4 °C with continuous stirring. Similarly, hexasomes, which lack an H2A/H2B dimer, were prepared by mixing the dimer, tetramer, and DNA in a 1:1:1 ratio (24). Both nucleosomes and hexasomes were purified using a 1 mL Source Q 4.6/100 column (Cytiva). Fractions containing the purified samples were pooled, dialyzed to 50 mM NaCl, concentrated using a Centricon with a 10-kDa cutoff (Millipore), and stored at 4 °C.

**Table 2.**
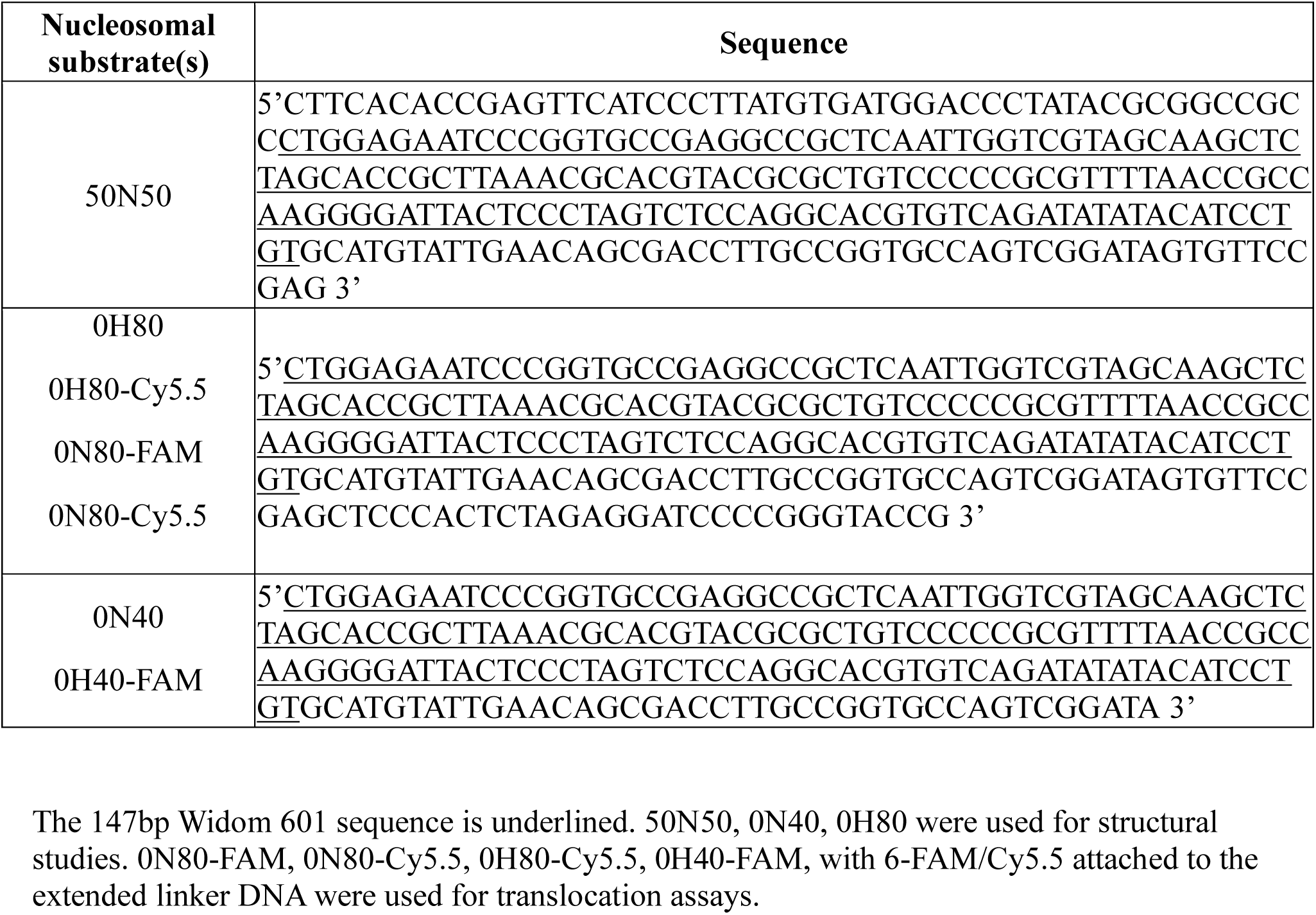
– DNA sequences of hexasomes and nucleosomes.

**Table 3.**
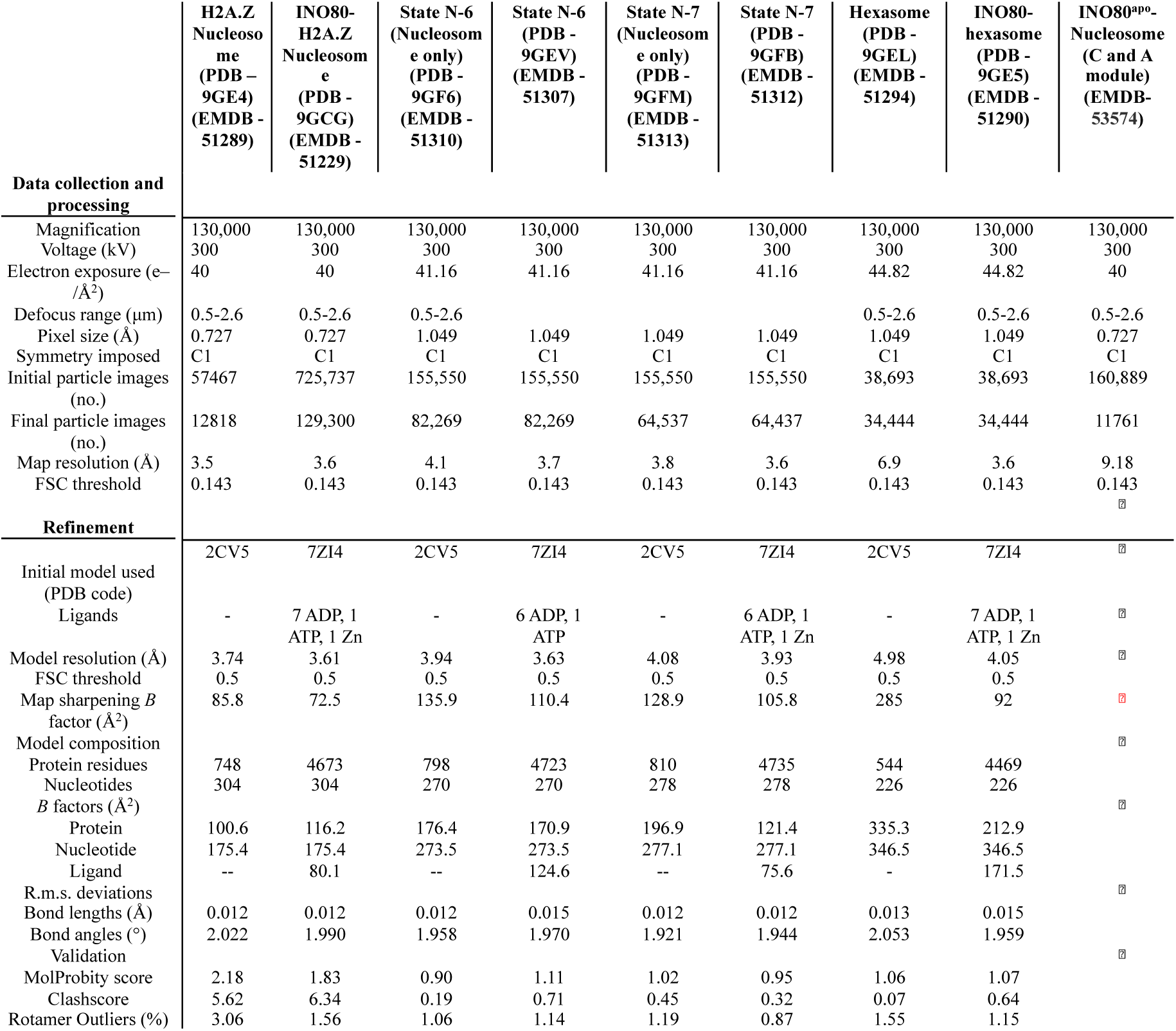
Cryo-EM data collection, refinement and validation statistics.

### Nucleosome and hexasome sliding assay

The translocation activity of INO80 on nucleosomes and hexasomes was measured as previously reported (24,29,31).Briefly, 6-Carboxyfluorescein labeled 0N80 nucleosomes and Cy5.5-labeled 0H80 hexasomes were used to determine the activity of INO80 and mutants. For the sliding assay, INO80 or its mutant at a concentration of 75 nM was mixed with 50 nM nucleosomes or hexasomes in a buffer containing 25 mM HEPES (pH 8.0), 60 mM KCl, 7% (v/v) glycerol, 0.1 mg/ml BSA, 2 mM MgCl_2_, and 0.25 mM DTT. The samples were pre-incubated at 37 °C for 2 minutes, and the sliding reaction was initiated by adding ATP to a final concentration of 1 mM. Samples were collected at various time points, including before ATP addition, at 30 seconds, and at 1, 2, 5, 10, 15, 30, and 60 minutes. Reactions were stopped by adding Lambda DNA (0.2 mg/ml; NEB). For loading, 1 µl of 50% (v/v) glycerol was added to each sample, and 6 µl of each sample was loaded onto a 3–12% acrylamide Bis-Tris native gel (Invitrogen). Gels were run at 110 V for 90 minutes. Visualization was performed using the Typhoon imaging system (GE Healthcare).

For hexasomes, the gels were run for a longer period—first at 110 V for 90 minutes, followed by an additional 30 minutes at 150 V to achieve better separation of translocated versus non-translocated hexasomes. All gels were then quantified using the ImageJ software.

To account for minor variations in sample loading and the presence of small amounts of nucleosomes in hexasome samples (and vice versa), a correction was applied. The percentage of the substrate’s band intensity (S%) was calculated at each time point relative to the combined intensity of the substrate, intermediate products, and final product. The S% value of the –ATP samples was used as a baseline, representing 100% non-remodelled substrate, and this baseline was then used to calculate the fraction of remodelled substrates. Graphs were generated using GraphPad Prism version 9.5.1 (Dotmatics) by fitting the data to a single-phase exponential decay model described by the following equation –

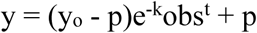

where *y*_0_ is the initial fraction product, *kobs* is the observed rate constant, and *p* is the fraction product at the plateau.

### ATPase assay

The ATPase activity of INO80 and mutants was determined using a NADH-coupled ATPase assay (31). INO80 or mutants (75 nM) were incubated in an assay buffer containing 25 mM HEPES (pH 8.0), 50 mM KCl, 1 mM DTT, 2 mM MgCl_2_, and 0.1 mg/ml BSA, along with 0.5 mM phosphoenolpyruvate, 1 mM ATP, 0.1 mM NADH, 25 U/ml lactate dehydrogenase, and pyruvate kinase (Sigma-Aldrich) at 37 °C in a final volume of 50 μl. The decrease in NADH concentration was monitored over 90 minutes using non-binding, 384-well black plates (Greiner Bio-One). Fluorescence was measured at an excitation wavelength of 340 nm and an emission wavelength of 460 nm using a Tecan Infinite M100 plate reader (Tecan). The ATPase rate was determined in the presence of 50 nM nucleosomes or hexasomes, with ATP turnover calculated using the maximal initial linear rates, corrected for a buffer blank, using the following equation –

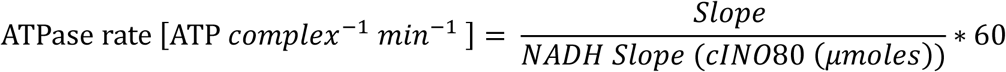

Where cINO80 is the concentration of human INO80 in micromoles, and NADH slope is the slope of the calibration curve calculated by carrying out titration of varying ADP concentrations ranging from 0 to 100 µM in 10 µM increments to constant NADH concentration. All the experiments were performed in triplicates.

### Competition assay

Nucleosome and hexasome translocation were simultaneously compared within the same reaction. For this, a mixture of 50 nM 6-Carboxyfluorescein (6-FAM) labeled 0N80 nucleosomes and 50 nM Cy5.5-labeled 0H40 hexasomes was incubated with 50 nM of either wild-type INO80 or mutant INO80. The competition assays were performed in the same manner as the individual translocation assays. Gels were run for 1.5 hours at 100 V, followed by an additional 30 minutes at 150 V, and then scanned for both 6-FAM and Cy5.5 signals. In parallel, the translocation of H2A.Z-containing 0N80 Cy5.5-labeled nucleosomes was compared to canonical nucleosomes under similar conditions, with gels run for 1.5 hours at 100 V. Quantification of the gels was carried out as described in the above section.

### Vitrification of the INO80-nucleosome/hexasome complex

For cryo-EM, freshly purified and highly concentrated INO80 complex was mixed with 0N40 nucleosomes or 0H80 hexasomes at final concentrations of 800 nM and 500 nM, respectively. The mixture was dialyzed in 2 L of dialysis buffer (20 mM HEPES, pH 8.0, 50 mM NaCl, and 0.25 mM TCEP) for 2 hours using Slide-a-lyzer dialysis tubes (Thermo Fisher Scientific).

ADP and MgCl_2_ were prepared at 10X concentration in a 2X Buffer (60 mM HEPES pH 8.0, 100 mM NaCl, 8 mM MgCl_2_, and 1 mM DTT). BeF_2_ and NaF were also prepared at 10X concentration in the same buffer. The dialyzed INO80-nucleosome complex or INO80-hexasome complex was then supplemented with ADP-MgCl_2_ and BeF_2_-NaF from these stock solutions to reach a final 1X concentration. This complex was incubated on ice for 30 minutes. β-Octyl glucoside (Roth, Germany) was added to a final concentration of 0.05%. The complex was vitrified on glow-discharged R2/1 copper mesh 200 grids (Quantifoil) by applying 4.5 μl of the sample, followed by a 2.2-second blot time using a Leica EM GP (Leica).

### Electron microscopy and data collection

Movies of INO80-nuclesosome embedded in vitreous ice were captured at liquid nitrogen temperature using a Titan Krios G3 transmission electron microscope (Thermo Fisher Scientific). This microscope was equipped with a K2 Summit direct electron detector (Gatan) and a BioQuantum LS Imaging Filter (Gatan). The movies were recorded in counting mode using EPU acquisition software (Thermo Fisher Scientific) at a magnification of 130,000X, resulting in a pixel size of 1.059 Å/pixel, with a nominal defocus range of –1.1 to 2.9 μm. Each movie received a total electron dose of approximately 40 to 46 e^−^/Å², distributed over 40 frames, with an exposure time of 250 ms per frame. Movies of INO80-Hexasome particles embedded in vitreous solution were collected at liquid nitrogen temperature using a Titan Krios G3 transmission electron microscope (Thermo Fisher Scientific) equipped with a Falcon 4 Direct Detection Camera and BioQuantum LS Imaging Filter (Gatan) operated at 300 kV acceleration voltage. The movies were recorded in counting mode using EPU acquisition software (Thermo Fisher Scientific) at 130,000X magnification with a pixel size of 0.727 Å/pixel and nominal defocus range of –1.1 to –2.6 μm. The total electron dosage of each movie was 40 e^−^/Å^2^, fractionated into 40 movie frames and exposure time of 3.19 seconds per frame.

### Cryo-EM data processing for INO80 complex

Movie frames were motion-corrected using MotionCor2 v1.4.5 (40). All subsequent cryo-EM data processing steps were performed using cryoSPARC v3.3.1 (41), with resolutions calculated using the gold-standard Fourier shell correlation criterion (FSC = 0.143). CTF parameters were estimated using patch CTF estimation (multi). The same data processing workflow was uniformly applied to all datasets used in this study, and data collection and refinement statistics are summarized in Supplementary Table 3.

A common feature of the INO80 complex across all analyzed states (H-3, N-7, N-6, NZ-7) is the presence of well-resolved structural elements (f**igs. (S1),(S2**),(**S3**),(**S4**). The RuvBL1/2 module is clearly resolved in all states, exhibiting an almost identical conformation (**figs. S1a, S2a, S3a, and S4a**). It forms a heterohexamer composed of RuvBL1 and RuvBL2, which serves as a scaffold for the assembly of additional subunits. The density for ADP is observed in the six nucleotide-binding pockets of RuvBL1/RuvBL2 in the maps of all states (**figs. S1b, S2b, S3b and S4b**).

The ARP5/IES6 subunits, located on the opposite side of the Ino80^motor^ (at different SHL positions depending on the state), are well resolved in all reconstructions (**figs. S1c, S2c, S3c and S4c**). The two RecA-like lobes of the ATPase domain (N-lobe and C-lobe) are interrupted by a large insertion, which is clearly threaded through the RuvBL1/2 hexamer, consistent with previous INO80–nucleosome structures from different species (**figs. S1d, S2d, S3d and S4d**) (28) (29)(31).

The Ino80^motor^ is well resolved in all states (**figs. S1e, S2e, S3e and S4e**), and its C-lobe interaction with nucleosomal/hexasomal DNA is clearly observed (**figs. S1f, S2f, S3f and S4f**). The IES2 subunit is also well resolved in all states (**figs. S1g, S2g, S3g and S4g**). Notably, additional map density corresponding to the N-terminal region of IES2 is visible in states N-7 and N-6. Focused refinement on IES2 and the nucleosome in these states further improved the quality of the maps and aided model interpretation (**figs. S2g, S3g**).

### Cryo-EM data processing for INO80-hexasome complex

For the INO80-Hexasome complex, initial particle picking was conducted using the blob picker. The selected particles were subjected to 2D classification. Defined 2D classes were chosen and used as input for a Topaz training job (42,43). The Topaz model was used for particle picking across 19,515 micrographs, yielding 981,468 particles, extracted with a box size of 440 pixels and a pixel size of 0.727 Å. Following 2D classification and ab initio reconstruction, the most well-defined classes were selected and subjected to heterogeneous refinement. The class with the best-defined features, comprising 34,444 particles, was chosen for further refinement, resulting in a final resolution of 3.6 Å for the INO80-Hexasome complex after non-uniform refinement (44).

To obtain a focused map around the hexasome, the volume was imported into ChimeraX 1.7.1(45). The “segment map” function in ChimeraX was used for splitting the volume. The first volume was generated for the hexasome, and the second volume was generated for RuvBL1/2, Ino80^motor^, ARP5/IES6 subunits, and IES2 (aa 208-342). The masks for hexasome (lowpass filter: 12, dilation radius: 15) as well as the RuvBL1/2, Ino80^motor^, ARP5/IES6 subunits, and IES2 (208–342) (lowpass filter: 12, dilation radius: 0) were then generated using these volumes in cryoSPARC v3.3.1 (41). Particle subtraction was thereafter done using the mask generated for RuvBL1/2, Ino80^motor^, ARP5/IES6 subunits, and IES2 (208–342). These subtracted particles were used against the mask generated for the hexasome to do the local refinement in cryoSPARC v3.3.1 (41) using the pose/shift Gaussian during alignment (standard deviation of prior over rotation: 10, standard deviation of prior over shifts: 5). Thus, a focused map was produced for the hexasome at a resolution of 6.9 Å.

In the INO80–hexasome complex, some 2D classes exhibited fuzzy map densities attached to the INO80-hexasome complex. These 2D classes (58,969 particles) were selected and re-extracted from the micrographs with a larger box size of 800 pixels. After another round of 2D classification, these particles underwent ab initio reconstruction followed by heterogeneous refinement. The resulting volume containing 26,073 particles, showing clear density for both the C and A-module, was further subjected to 3D classification. This classification further sorted the particles, isolating a subset of 22,112 particles that displayed density for both the C and A-module together. Subsequent non-uniform refinement produced a low-resolution map at 8.95 Å. The map density for the C module appeared clearer compared to the A-module due to the flexible nature of the latter. To enhance the quality of the flexible A-module, particles were further processed using 3D-flex refinement (46) with default parameters. The final map obtained from this 3D-flex refinement was then used for docking the previously determined C module structure and the AlphaFold3-predicted (47) structure into the corresponding map densities. The processing scheme is illustrated in **supplementary fig 5.**

### Cryo-EM data processing for INO80-nucleosome complex

The initial processing for INO80-nucleosome dataset was processed similarly as mentioned above. After topaz training (42,43), the resulting Topaz model was then applied as a template for particle picking across 19,528 micrographs, yielding 546,516 particles, extracted with a box size of 440 pixels and a pixel size of 1.046 Å. After selecting 2D classes with defined features, a round of ab initio reconstruction with five classes was performed. Classes with most well-defined features were chosen for heterogeneous refinement into three classes, followed by 3D classification, resulting in two distinct classes, named state N-6 and state N-7. Both classes were further refined using non-uniform refinement (44), achieving final resolutions of 3.7 Å for state N-6 and 3.6 Å for state N-7.

To obtain focused maps around the nucleosome and IES2 (residues 140-189) in both states N-6 and N-7, the volumes were imported into ChimeraX 1.7.1. The “segment map” function in ChimeraX1.7.1(45) was used for splitting the volumes for both states (N-6 and N-7). The first volume was generated for the nucleosome and IES2 (140–189), and the second volume was generated for RuvBL1/2, Ino80^motor^, ARP5/IES6 subunits, and IES2 (208–342). The masks for nucleosome and IES2 (140–189) (lowpass filter: 12, dilation radius: 15) as well as the RuvBL1/2, Ino80^motor^, ARP5/IES6 subunits, and IES2 (208–342) (lowpass filter: 12, dilation radius: 0) were then generated using these volumes in cryoSPARC v3.3.1 (41). Particle subtraction was thereafter done using the mask generated for RuvBL1/2, Ino80^motor^, ARP5/IES6 subunits, and IES2 (208–342). These subtracted particles were used against the mask generated for the nucleosome and IES2 (140–189) to do the local refinement in cryoSPARC v3.3.1 (41) using the pose/shift Gaussian during alignment (standard deviation of prior over rotation: 10, standard deviation of prior over shifts: 5). Thus, focused maps were produced at the resolutions of 3.72 Å for state N-6 and 3.80 Å for state N-7, respectively. The processing scheme is illustrated in **supplementary fig 6.**

### Cryo-EM data processing for INO80-H2A.Z nucleosome complex

The INO80 H2A.Z–nucleosome complex was processed similarly as mentioned above. Briefly, from a total of 36,316 micrographs, initial processing steps described above were performed. Particle picking was carried out using a blob picker, followed by particle extraction from micrographs with a box size of 400 pixels and a pixel size of 1.046 Å, yielding 299,459 particles. These particles underwent multiple rounds of 2D classification. After final 2D classification, 5,076 particles from well-defined classes were selected and used as input for training a Topaz model. This trained Topaz (42,43) model was subsequently employed for particle picking across all 36,316 micrographs, resulting in 3,108,522 particles, extracted with a box size of 440 pixels. After additional rounds of 2D classification, particles displaying clearly defined features were selected. These particles underwent ab initio reconstruction into six classes. The best-defined classes were selected and subjected to heterogeneous refinement into six classes, followed by 3D classification, which yielded two similar classes. These two classes, comprising 94,892 particles, were combined and further refined using non-uniform refinement (44), achieving a final resolution of 3.6 Å.

To obtain a focused map around the H2A.Z–nucleosome, the volume was imported into ChimeraX 1.7.1. The “segment map” function in ChimeraX was used for splitting the volume. The first volume was generated for the nucleosome, and the second volume was generated for RuvBL1/2, Ino80^motor^, ARP5/IES6 subunits, and IES2 (208–342). The masks for nucleosome (lowpass filter: 12, dilation radius: 15) as well as the RuvBL1/2, Ino80^motor^, ARP5/IES6 subunits, and IES2 (208–342) (lowpass filter: 12, dilation radius: 0) were then generated using these volumes in cryoSPARC v3.3.1 (41). Particle subtraction was thereafter done using the mask generated for RuvBL1/2, Ino80^motor^, ARP5/IES6 subunits, and IES2 (208–342). These subtracted particles were used against the mask generated for the nucleosome to do the local refinement in cryoSPARC v3.3.1 (41) using the pose/shift Gaussian during alignment (standard deviation of prior over rotation: 10, standard deviation of prior over shifts: 5). Thus, a focused map was produced for the nucleosome at a resolution of 3.5 Å.

Additionally, similar to observations in the INO80–hexasome complex, some particles displaying fuzzy density in the 2D classification were identified. These particles were selected and re-extracted with a larger box size of 800 pixels, followed by additional rounds of 2D classification. The well-defined 2D classes obtained were selected and used to train another Topaz model. This trained model was then applied for particle picking across the entire set of 36,316 micrographs, yielding 179,796 particles, extracted with a box size of 440 pixels. After multiple rounds of 2D classification, particles exhibiting clear features were chosen. These selected particles underwent ab initio reconstruction into two classes, and the class with the most clearly defined features was subjected to heterogeneous refinement into two classes. The better-defined class containing 11,540 particles was further refined using non-uniform refinement, resulting in a low-resolution map of 11.66 Å. Similar to the INO80–hexasome complex, the A-module map quality was poorer compared to the C module due to inherent flexibility. To improve the quality of the flexible A-module map, the particles were further processed using 3D-flex refinement (46) with default parameters. The final map obtained from this 3D-flex refinement was subsequently used for docking the previously determined C-module structure along with the AlphaFold 3-predicted structure into their corresponding map densities. The processing scheme is illustrated in **supplementary fig 7**.

### Cryo-EM data processing for INO80^apo^-nucleosome complex

For the INO80–nucleosome complex without nucleotide, a total of 30,012 micrographs were collected, and initial processing steps were performed as described above. Following multiple rounds of 2D classification, particles with well-defined features were selected and used to train a new Topaz model (42,43). This trained model was then applied to the full dataset, yielding 605,639 particles, which were extracted with a box size of 440 pixels.

As in previous datasets, several 2D classes showed fuzzy densities, suggestive of partially resolved regions. These particles were processed using a similar workflow. Heterogeneous refinement and subsequent 3D classification yielded a reconstruction with clearly defined C and A-module densities. This reconstruction was further refined using non-uniform refinement (44), resulting in a final resolution of 9.18 Å from 16,786 particles.

Due to the intrinsic flexibility of the A-module, its density remained suboptimal, prompting the use of 3D-flex refinement to improve map quality. The final 3D reconstructions for both classes were used to dock the previously determined C-module structure and the AlphaFold-predicted A-module structure into their respective densities. The processing workflow is summarized in **supplementary fig 8.**

### Model building and refinement

The model for the INO80 C-module-nucleosome complex was initially constructed for state N-7 and then used as the basis for the model building of state N-6, INO80-H2A.Z containing 50N50 nucleosomes, as well as for the INO80-hexasome complex state H-3. These models were built into B-factor sharpened maps derived from focused refinements. The starting models were obtained from PDB entries 7ZI4 (for the C-module) (28), and 2CV5 (48) (for the nucleosome). These models were first rigid-body fitted using ChimeraX version 1.7.1 (45) and then model rebuilding was performed using COOT version 0.9.8.93 (49). For State N-7, DNA was modeled beyond SHL-3 and fitted into the density using German-McClure restraints in COOT. The N-terminal region of IES2 (140–189), in both states N-6 and N-7 was modeled by rigidly fitting the AlphaFold–predicted structure of IES2 (AF-Q9C086-F1-v4) (50) into the corresponding cryo-EM densities, and refined using COOT. The Model building and refinement were carried out iteratively using interactive molecular dynamics flexible fitting in ISOLDE v1.4 (51). Reciprocal space refinement, using jelly-body restraints, was performed with REFMAC5 (52) against maximum-likelihood weighted structure factors calculated from cryo-EM half-maps. Additional model building was conducted in COOT against the maximum-likelihood estimate of the expected true map calculated by REFMAC5. Final model corrections were made using ISOLDE against the same REFMAC5 map, followed by a concluding round of reciprocal space refinement with jelly-body restraints in REFMAC5.

All structural figures were prepared using UCSF ChimeraX 1.7.1(45).

## Results

### Human INO80 can slide both hexasomes and nucleosomes

We recombinantly reconstituted the core *Homo sapiens* (*Hs*) INO80 complex, consisting of an evolutionarily conserved C-(Ino80^motor^, ARP5, IES6, IES2, RuvBL1, RuvBL2) and A-module (ARP8, Actin, ARP4, YY1) (**figs S9a-c**). This complex follows prior work, however with the addition of the transcriptional regulator YY1(28). In the presence of ATP, the human INO80 complex demonstrated robust nucleosome translocation activity, similar to what has been previously observed with *Hs,*fungal and yeast INO80 complexes (23,28,29). We next tested hexasome sliding activity of human INO80 and found robust sliding capability (**figs. 1a-c)** We used 80 bp of extranucleosomal DNA for nucleosome and 40 bp extranucleosomal DNA for hexasome substrates, to minimize influences of linker DNA lengths on sliding rates caused by additional nucleosomal DNA unwrapping in the case of the hexasome. Our results show that human INO80, like yeast and fungal INO80, can efficiently slide hexasomes (23)(24). Notably, we do not see a strong preference for either hexasomes or nucleosomes if sufficient extranucleosomal DNA (∼40 bp) is available (23)(24) (**fig S9d**).

### Structure of Human INO80 bound to hexasomes

To understand how INO80 recognizes hexasomes, we determined structures in complex with hexasome (0H80) using single-particle cryo-electron microscopy (cryo-EM). The structures were determined in the presence of the ATP analog ADP-BeF_3_ without any chemical crosslinking.

We determined the structure of INO80-0H80 at 3.6 Å, with a focused map of the hexasome at 6.9 Å. The map revealed the INO80 C-module, while the A-module was not visible due to flexibility. The INO80 C-module engages the “proximal face” of the hexasome i.e., the side where the H2A/H2B dimer is missing. Due to the missing H2A/H2B dimer, an additional ∼46 bps of DNA were unwrapped from the histone core. The Ino80^motor^ domain binds DNA at the resulting new entry position of linker DNA, which is superhelical location –3 (SHL-3). Similar to yeast and fungal INO80, human INO80 undergoes a ∼145° “rigid body” spin rotation around the “axle” of the hexasome “wheel” in relation to its location on the nucleosome. As a consequence, the ARP5/IES6 subunits binds DNA on the opposite side of the hexasome near the dyad (SHL0) and SHL+1. We denote this INO80 binding state “H-3” with the Ino80^motor^ domain located at <u>h</u>exasome SHL<u>-3</u> (23)(24) **(fig 1d)**.

**Figure 1:**
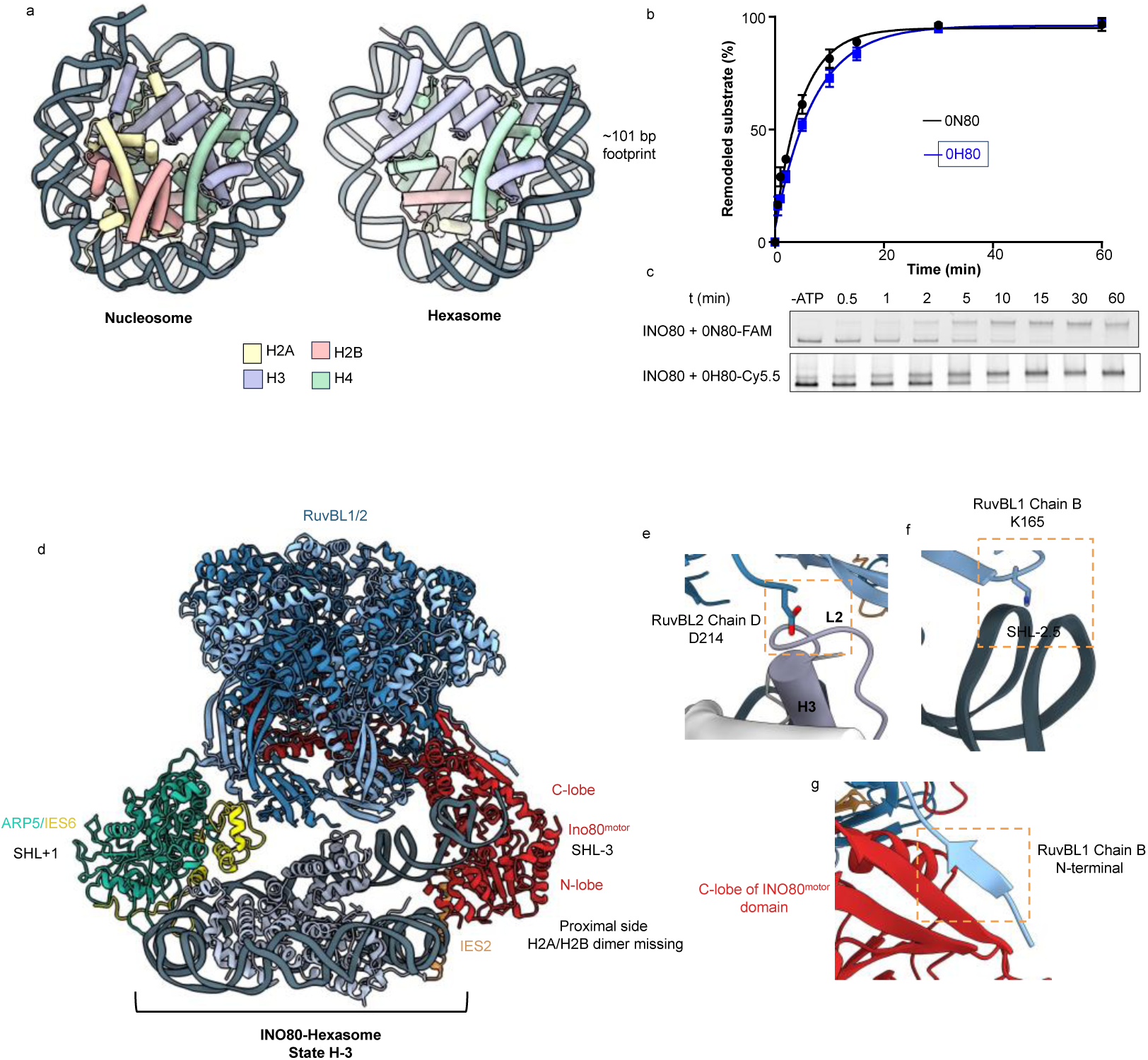
Structure of INO80 bound to Hexasome. **(a)** Cartoon depiction of a canonical nucleosome (PDB: 2CV5) (48) and the hexasome from this study. The hexasome was reconstituted with Widom 601 DNA sequence (39) (80 bp linker on one side), spanning a 101 bp footprint and lacking the H2A-H2B dimer. **(b)** Translocation activity of INO80 on nucleosome (0N80) and hexasome (0H80) substrates is shown (n=3). Fraction of centrally positioned nucleosome or hexasome substrate is indicated as remodeled substrate in percent. **(c)** Native PAGE analysis of sliding of end positioned nucleosomes and hexasomes by INO80 is shown. **(d)** Cryo-EM structure of the INO80 C-module bound to the hexasome. INO80 binds on the proximal side of the hexasome which lacks the H2A-H2B dimer (state H-3). (**e**) Close-up view of the interaction between INO80 and the histone core within the hexasome. Residue D214 in chain D of RuvBL2 is located within 5 Å of residues in the α2 helix of histone H3. **(f)** Additionally, residue K165 in chain B of RuvBL1 interacts with the phosphate backbone of the unwrapped DNA near SHL-2.5. (**g**) The N-terminus of chain B of RuvBL1 forms an additional strand on the edge of a β-sheet of the C-terminal lobe of the motor domain.

Hexasome recognition is mostly governed through binding of hexasomal DNA, with the exception of one RuvBL2 aspartic acid residue (D214) contacting the H3 loop 2. This interaction might stabilize this H3/H4 pair at the entry side DNA **(fig 1e)**. The predominantly DNA mediated hexasome binding contrasts yeast and fungal orthologs, where the Arp5 grappler insertion (largely missing in human INO80) binds either the H2A/H2B acidic patch (nucleosome) or H3/H4 (hexasome). DNA mediated contacts are found at the ATPase (entry DNA), where the RuvBL1 K165 residue binds the phosphate backbone of hexasomal DNA near the SHL-2.5 position and may stabilize hexasomal DNA at the new linker DNA entry site **(fig 1f)**. Interestingly, residues 3-IEEV-6 of the same RuvBL1 protomer form an additional β-strand attached to the β-sheet of the Ino80^motor^ C-lobe **(fig 1g).** This type of β-strand addition has not been observed in the fungal and yeast INO80 complexes and possibly rigid the interaction of the Ino80^motor^ with the RvBL1/2 domain.

### INO80 binds nucleosomes in different positions

We subsequently determined the structure of human INO80 bound to nucleosomes (0N40) and observed two distinct nucleosome binding states, denoted state N-7 and state N-6, from the same dataset **(figs 2a,b).** In state N-7 (3.6 Å, with the nucleosome locally refined to 3.8 Å), the Ino80^motor^ binds nucleosomal DNA at the SHL-7 position, while ARP5/IES6 subunits are located at SHL-3 (**fig 2a**). This position is similar to the previously determined structure of human INO80 in complex with the 50N25 nucleosome (28). Interestingly, we observed with N-6 a second state (3.7 Å, with the nucleosome locally refined to 4.1 Å), where human INO80 underwent a spin rotation by one SHL, resulting in the ATPase domain binding at SHL-6 and the ARP5/IES6 module at SHL-2 (**fig 2b**). This position is similar to nucleosome binding of yeast and fungal INO80 (23,29). Transitioning from state N-7 to state N-6 involves detachment of DNA from H2A at SHL-6, leading to unwrapping of ∼10 bp from the histone core (**fig 2c**). In addition, the H3 tail moves away from the Ino80^motor^ (**figs 2d,e**) and DNA contacts made by the proximal-side histone H3 αN and H2A loop 2 are broken, resulting in the partial exposure of the H2A/H2B dimer (**figs S10a,b**).

**Figure 2:**
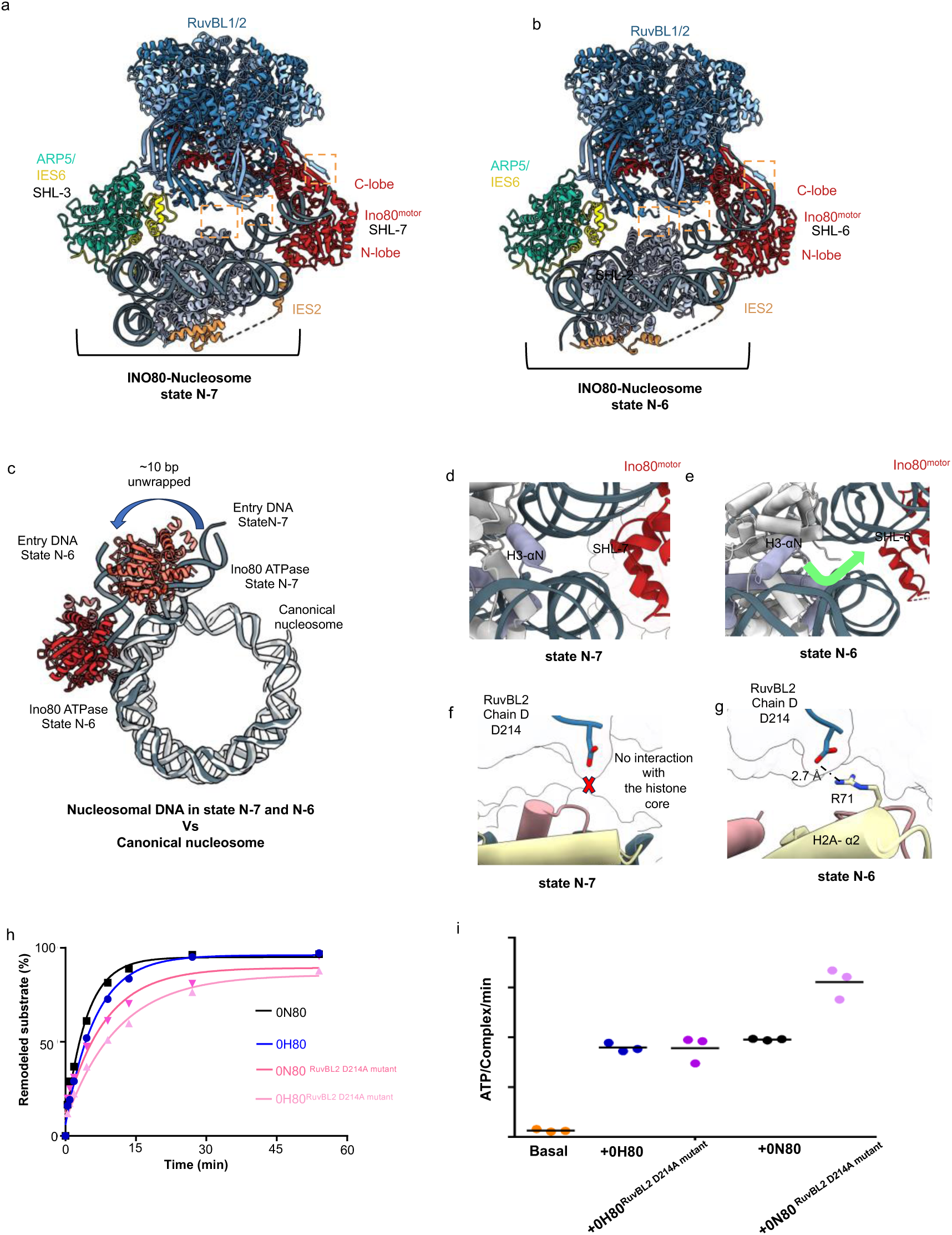
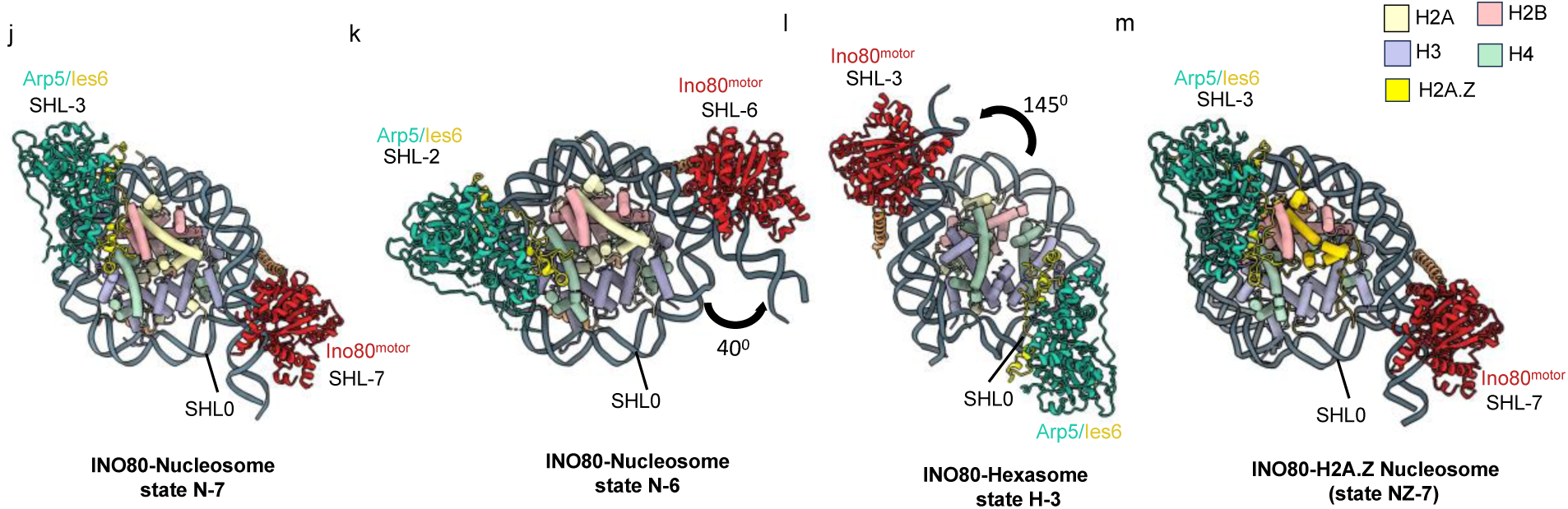
INO80 binds nucleosomes in different positions. (**a,b**) Cryo-EM structures of INO80 core module bound to nucleosomes in (a) states N-7 and (b) N-6. (**c**) Structural comparison of nucleosome states N-7 (in gray) and N-6 (in light gray) relative to the canonical nucleosome (in white) (PDB: 2CV5) (66). The transition from state N-7 to state N-6 results in unwrapping of approximately 10 bp from the histone core. The histone core is aligned. (**d**) The N-terminal H3 tail of the nucleosome in state N-7 is positioned near the Ino80^motor^ C-lobe. Despite its proximity, no density map signal was observed to indicate a direct interaction between the H3 tail and the ATPase subunit. (**e**) In state N-6, the H3 tail moves further away from the Ino80^motor^ C-lobe, due to the translocation of INO80 **(f**) No interactions were observed between histone core residues and RuvBL2 in state N-7. (**g**) In contrast to state N-7, where no interaction was observed with the histone core, state N-6 shows a strong salt bridge interaction between residues R71 of histone H2A (α2) and D214 of the RuvBL2 subunit of INO80. (**h**) Evaluation of the remodeling activity of INO80 RuvBL2 D214A mutants. Band intensities of remodeled and unremodeled nucleosome/hexasome species were quantified, and the fraction of remodeled nucleosomes/hexasomes was plotted against time. Data points were fitted using an exponential equation (see method). Mean values and individual data points (n = 3, technical replicates) are shown. (**i**) ATPase activity of INO80 RuvBL2 D214A mutants, with and without stimulation by nucleosomes/hexasomes. ATPase rates were calculated from the linear region of the raw data and corrected using a buffer blank. Mean values and individual data points (n = 3, technical replicates) are presented. (**j-m**) Comparison of INO80 structures bound to different substrates. Shown are the multiple states of the INO80–nucleosome complex (N-7, N-6, and INO80–H2A.Z nucleosome (NZ-7), centrally positioned (50N50) alongside the INO80–hexasome structure (H-3). Superhelical location (SHL) is defined by DNA translocation, with regions proximal to the DNA entry site designated as “-” and those further along the DNA path designated as “+”. In state N-7, the Ino80^motor^ binds to the nucleosome at SHL-7, and the ARP5/IES6 subunits bind at SHL-3. In state N-6, the Ino80^motor^ and ARP5/IES6 shift to SHL-6 and SHL-2, respectively, with ARP5/IES6 rotating approximately 20 degrees compared to state N-7. In state H-3, the Ino80^motor^ binds to entry DNA at SHL-3, while ARP5/IES6 subunits bind near the dyad (SHL-0 postion). Notably, INO80 interacts with the hexasome from the side lacking the H2A/H2B dimer, undergoing an approximately 145-degree rotation relative to state N-7. Finally, in the centrally positioned nucleosome (50N50), INO80 binds similarly to state N-7.

RuvBL2 D214 forms a salt bridge with the H2A α2 residue R71 in N-6 but not N-7 (**fig 2f,g**). We hypothesize that D214 might help stabilize the histone core in both hexasomes and nucleosomes when DNA is unwrapped and that the N-6 state is the “active” state on nucleosomes. Consistently, INO80^RuvBL2_D214A^ showed decreased translocation activity for both nucleosomes and hexasomes compared to the wild type (**fig 2h**) (**figs S10c**). Hereby, the ATPase activity of INO80 was either unaffected (hexasome), or even slightly increased (nucleosome) compared to wild type (**fig-2i**), showing the reduction is not due to compromised ATP hydrolysis rate.

Given the distinct modes of nucleosome binding by INO80, we also investigated how human INO80 binds a nucleosome containing the H2A variant H2A.Z. H2A.Z possesses additional acidic residues particularly in the acidic patch region (19) (**figs S11a,b**). As previously observed, we also found that H2A.Z-containing nucleosomes are preferred substrates over canonical nucleosomes (29) (**fig S11c**). We determined the structure of human INO80 bound to a centrally positioned 50N50 H2A.Z-containing nucleosome using cryo-EM in the presence of ADP-BeF_3_ (State NZ-7). The overall resolution achieved was 3.5 Å, with a focused map resolution around the nucleosome of 3.6 Å. Despite the extended acidic patch on the H2A.Z-containing nucleosome, the binding mode of INO80 was similar to that of canonical nucleosomes in state N-7 **(fig S11d)**.

Comparing H-3, N-6 and N-7 (H2A– and H2A.Z-nucleosomes) states shows that human INO80 moves in a rigid body manner without substantial changes in the C-module structure (**figs 2j-m**). Since the structures for states N-7, N-6, H-3 and the H2A.Z nucleosome were all done in the presence of ADP-BeF_3_ the different nucleosome interaction modes are not a consequence of the nucleotide binding to the Ino80^motor^. The lack of histone contacts in conjunction with the spin rotation suggests that a defining feature of nucleosome/hexasome binding by human INO80 in respect to the point of entry DNA, a topological feature (see discussion). In summary, we reveal an intrinsic flexibility in nucleosome binding of human INO80 that also resolves the discrepancy in nucleosome recognition between human and fungal INO80 in earlier work.

### Both ARP5 and IES6 nucleosomal DNA binding are required for robust sliding

To further understand the remodelling of nucleosomes and hexasomes, we investigated the role of the ARP5/IES6 module. ARP5’s DNA-binding domain (DBD) (29) interacts with nucleosomal DNA in the minor groove at SHL positions –3 in state N-7 and –2 in state N-6, while it binds to hexasomal DNA at SHL position +1 in state H-3 (**fig 3a**). Previous studies with fungal INO80 have shown that the conserved DBD loop residues of ARP5 are essential for the sliding activity of fungal INO80 on both nucleosomes and hexasomes (24,29). However, we observed that also a loop of the IES6 subunit (residues H129-K142), which contains multiple positively charged residues, interacts with the DNA phosphate backbone in the major groove at SHL positions –4 in state N-7, –3 in state N-6, and +2 in the case of state H-3 (**fig 3a**). The multiple sequence alignment of the lES6 region (H129-K142) showed that the positively charged residues are partially conserved among species **(fig 3b)**. To test the functional relevance of the IES6 DNA binding, we mutated the DNA binding IES6 loop (G135A/K136A/K137A). These residues interact with the DNA phosphate backbone and their mutation significantly affected the sliding activity of INO80 while still maintaining robust ATPase activity for both nucleosome and hexasome substrates (**figs 3c,d) (fig S12a**). The interactions of IES6 with the histone core and its binding to nucleosomal/hexasomal DNA suggest that the IES6 loop region plays a regulatory role in human INO80 sliding.

**Figure 3:**
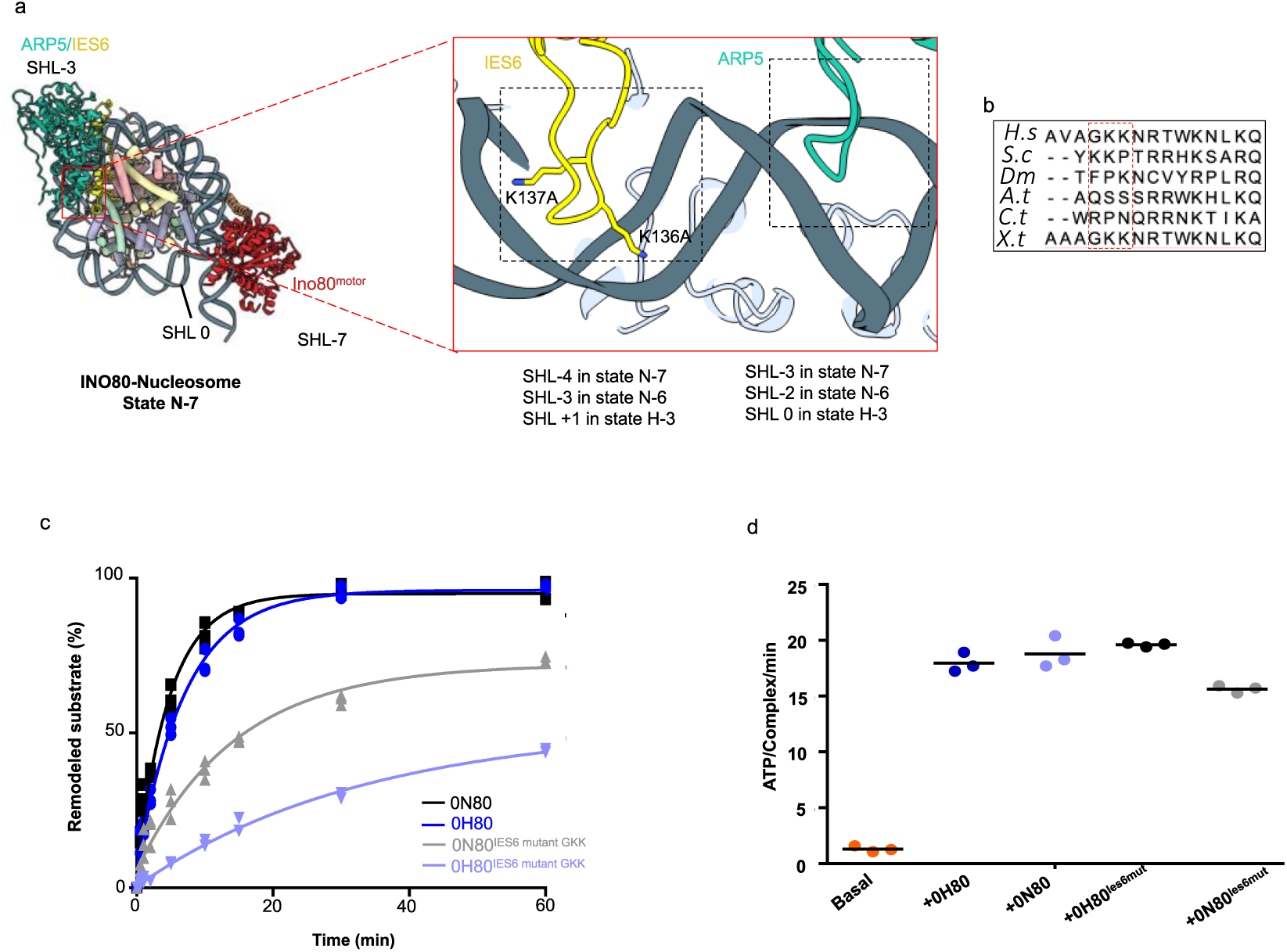
Both ARP5 and IES6 nucleosomal DNA binding are required for robust sliding. (**a**)Zoomed-in view of the interaction between the ARP5-IES6 subunits and nucleosomal/hexasomal DNA. ARP5 interacts with the phosphate backbone at the minor groove of DNA. This interaction occurs at SHL-3 in state N-7 and SHL-2 in state N-6. In state H-3, ARP5 interacts near the dyad. IES6 establishes additional contacts at the major groove of DNA, binding at SHL-4 in state N-7 and SHL-3 in state N-6 via its loop residues GKK (135–137). The IES6 loop residues GKK (135–137) interact with hexasomal DNA near the SHL +1 position. The INO80 ARP5/IES6 subunits show a similar interaction with the centrally positioned nucleosome as observed in state N-7. (**b**) Multiple sequence alignment is shown for the loop residues of IES6 involved in interactions with nucleosomal and hexasomal DNA. *H*.*s*., *Homo sapiens; S*.*c*., *Saccharomyces cerevisiae*; *D*.*m*., *Drosophila melanogaster*; *A*.*t*., *Arabidopsis thaliana*; *C*.*t*., *Chaetomium thermophilum*; *X*.*t*., *Xenopus tropicalis*. (**c**) Evaluation of the remodeling activity of INO80 IES6 GKK mutants. Band intensities of remodeled and unremodeled nucleosome/hexasome species were quantified, and the fraction of remodeled nucleosomes/hexasomes was plotted against time. Data points were fitted using an exponential equation (see method). Mean values and individual data points (n = 3, technical replicates) are shown. (**d**) ATPase activity of INO80 IES6 mutants, with and without stimulation by nucleosomes/hexasomes. ATPase rates were calculated from the linear region of the raw data and corrected using a buffer blank. Mean values and individual data points (n = 3, technical replicates) are presented.

### Acidic patch binding is required for nucleosome but not hexasome sliding

The H2A/H2B acidic patch is the most commonly recognized epitope on the nucleosome and has been shown to regulate the activity of various remodellers (53). Fungal ARP5 has an insertion domain (“grappler”) which directly binds the acidic patch via a foot region (29). Human ARP5 has a much smaller insertion domain that particularly lacks the acidic patch binding foot. However, human INO80 possesses the putative acidic patch binding by IES2. To examine the effect of the acidic patch on the INO80 chromatin remodeller, we mutated the H2A acidic patch residues (E61A/E64A/D72A/D90A), referred to as APM (**fig S12b**). The INO80 complex exhibited reduced sliding activity but retained robust ATPase activity with the APM (**figs S12c-d**). These data contrast findings for fungal INO80, where mutations of the acidic patch residues completely abolished nucleosome sliding activity (29). Interestingly, the APM had no effect in the case of hexasomes, suggesting that IES2 mediated acid patch contacts are important in the context of nucleosome remodelling but not hexasome remodelling (**figs.S4c-e**).

### IES2 functionally interacts with the acidic patch on nucleosomes but not hexasomes

In our INO80 structure with 0N40 nucleosomes, in both state N-6 and state N-7, we were able to improve the resolution of the IES2 subunit compared to the previously published structure (28), revealing additional density at the distal side of the nucleosome. Based on the shape, chemical environment, and AlphaFold2 modelling (AF-Q9C086-F1-v4) (50), this density corresponds to the N-terminus of IES2 (G140-S189). We built the model for this region of IES2 and found that it forms multiple contacts with the distal side histone and nucleosomal DNA. The resulting model includes IES2 residues G140-S189 (region I), T236-N263 (region II), and P271-R342 (region III), which make contacts with the histone core as well as with the Ino80^motor^ and with the RuvBL1/2 subunits of the INO80 complex. **(fig S13a)**. The N-terminal α-helix (G140-K158) of IES2 region I is positioned near the DNA exit site at SHL +5 in both state N-7 and N-6 (**fig 4a)**. We observed no density for DNA beyond SHL +5 in either state. When aligning the canonical nucleosome with the map density, the IES2 α-helix (G140-K158) causes steric hindrance at the DNA exit, suggesting that this interaction needs partially unwrapped exit DNA in a chromatin environment **(fig 4b)**. DNA unwrapping from both entry and exit sides has been reported in other chromatin remodellers like Chd1 and SWR1(6,54).

**Figure 4:**
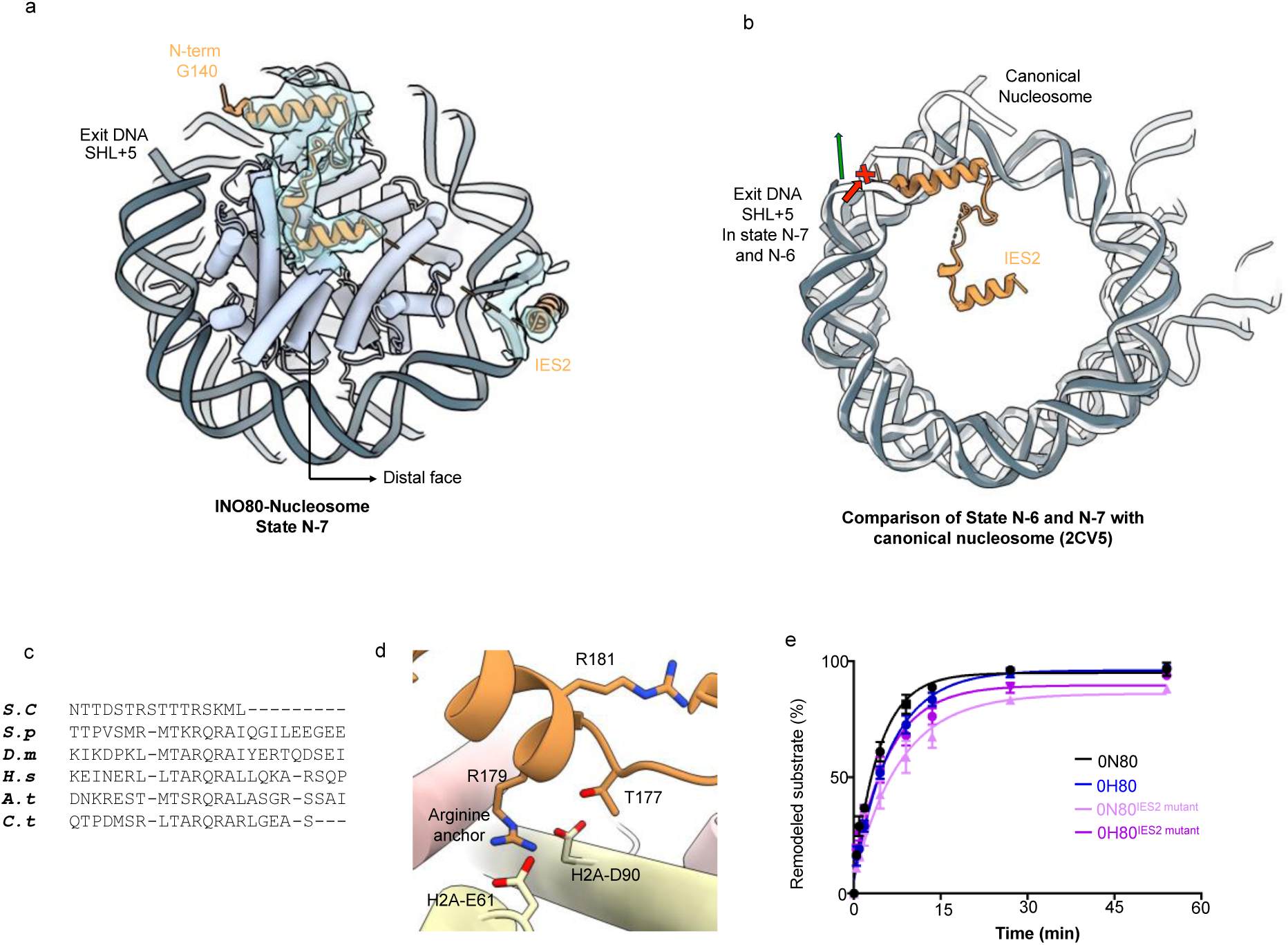
Roles of IES2: DNA Unwrapping from the Distal Side and its Engagement with the Acidic Patch. **(a**) The density map shows the modeled IES2 N-terminal helix (G140-S189) interacting with the distal side of the nucleosome in state N-7. (**b**) Comparison of IES2 interaction with nucleosomal DNA in states N-7 and N-6 with that of the canonical nucleosome (PDB: 2CV5) reveals that nucleosomal DNA in states N-7 and N-6 cannot follow the same path as in the canonical nucleosome. If it did, a steric clash would occur between the exit-side nucleosomal DNA and the IES2 N-terminal helix. The histone core is aligned (**c**) Multiple sequence alignment of IES2 subunit across species highlights the conserved arginine anchor residues (highlighted in light blue color), and partially conserved arginine residues (highlighted in light green color). *H*.*s*., *Homo sapiens; S*.*c*., *Saccharomyces cerevisiae*; *D*.*m*., *Drosophila melanogaster*; *A*.*t*., *Arabidopsis thaliana*; *C*.*t*., *Chaetomium thermophilum*; *X*.*t*., *Xenopus tropicalis*. (**d**) Interaction of conserved arginine residue (R179) of IES2 with the acidic patch residues (E61, D90) of histone H2A. (**e**). Evaluation of the remodeling activity of INO80 IES2 mutants. Band intensities of remodeled and unremodeled nucleosome/hexasome species were quantified, and the fraction of remodeled nucleosomes/hexasomes was plotted against time. Data points were fitted using an exponential equation. Mean values and individual data points (n = 3, technical replicates) are shown.

**Figure 5:**
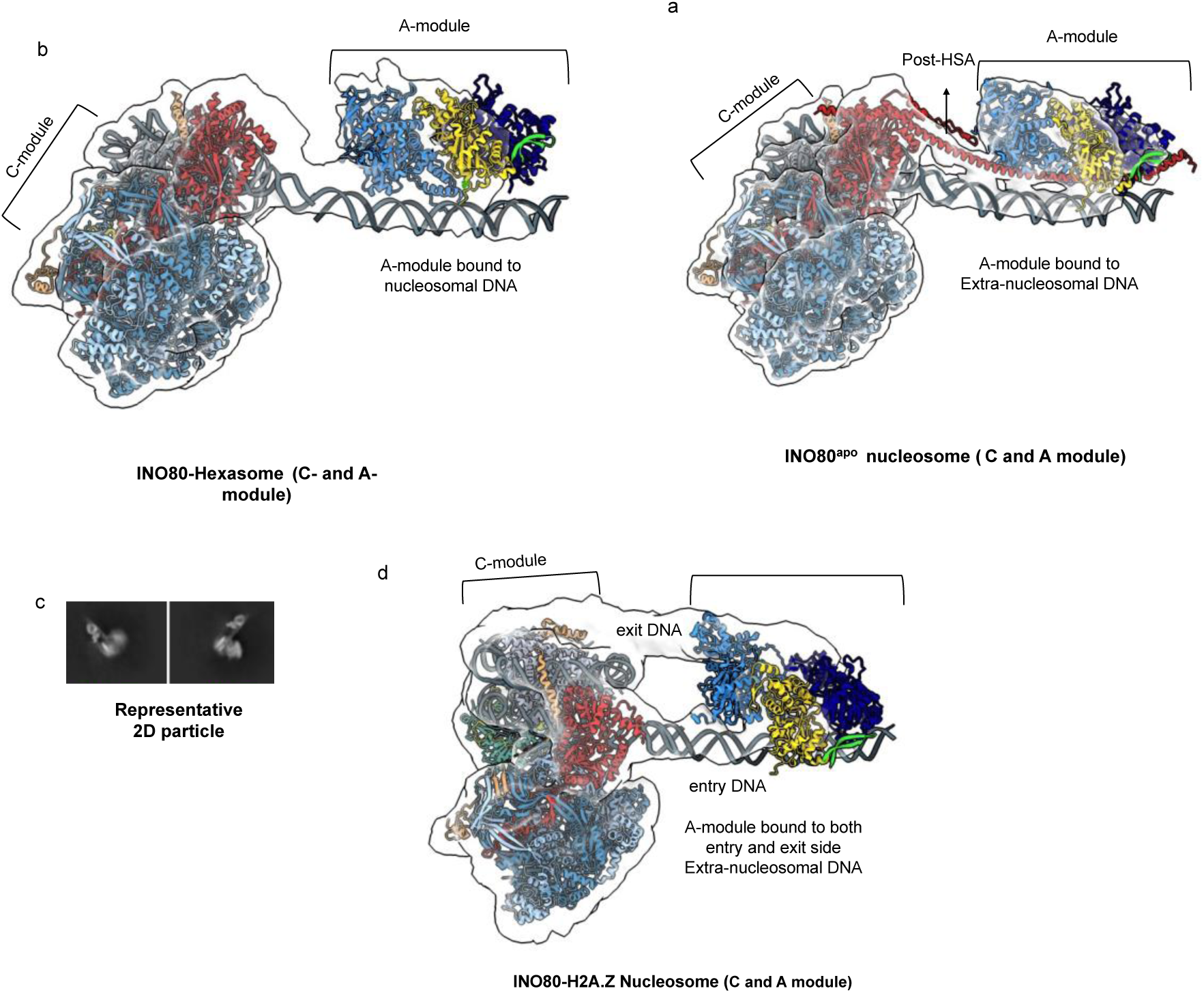
Overall structure of the INO80 core and A-module bound to nucleosome/ hexasome. **(a,b)** Maps generated by 3D flexible refinement (46) showing the INO80 core module combined with the AlphaFold-predicted A-module structure fitted into the density for (a) INO80^apo^–nucleosome (core and A-module), and (b) INO80–hexasome (core and A-module). Map density for the post-HSA region is clearly visible in the INO80^apo^–nucleosome map but absent in the INO80–hexasome reconstruction. (**c**) Representative 2D particle classes used for reconstruction of the core-plus-A-module complex, illustrating A-module binding to both the entry and exit sides of extranucleosomal DNA. (**d**) Map from 3D flexible refinement of the core-plus-A-module complex highlighting density at the entry and exit sites of extranucleosomal DNA. Map density for the post-HSA region is not observed.

Region I also exhibits a highly conserved RxR arginine anchor motif, which binds the acidic patch (**fig 4c,d**). Mutating R179 and R181 to alanines decreased sliding activity (**fig 4e**) (**fig S13b**), but increased the ATPase activity of INO80 in the nucleosome context three-fold (**fig S13c**). A previous INO80 delta IES2 construct was deficient both in nucleosome sliding and ATPase activity (37). Finally, no effect of the RxR mutation was observed on hexasome sliding activity or hexasome-stimulated ATPase activity, consistent with a lack of density for IES2 at the hexasome distal histone surface including the acidic patch.

The region II of IES2 contains the throttle helix which interacts with the nucleosomal DNA near the major groove at different SHL positions in state N-6, N-7 and H-3 (**fig S13a**). Region III of IES2 binds RuvBL2. Both regions essentially bind as previously established (37,55).

In summary, we can conclude that the throttle helix is important for ATPase activity while the acid patch anchor RxR is critical to convert ATPase activity into sliding of nucleosomes, while it does not appear to play a role in mobilizing and binding to hexasomes.

### Overall structure of the INO80 C and A-module bound to nucleosome/ hexasome

To define the overall architecture of the C and A-module, we determined the structure of INO80 in the absence of nucleotide, as previously done for fungal INO80 under nucleotide-free conditions (31). 2D classification revealed particles containing the INO80 C-module along with density corresponding to the A-module. To improve resolution, these particles were re-extracted using a larger box size (*see method section*).

Applying the same strategy to the other datasets (INO80–hexasome, INO80–nucleosome, and INO80–H2A.Z) revealed some 2D classes with A-module density in the INO80–hexasome and INO80–H2A.Z dataset (*see method section and supplementary figure*). However, no additional A-module density was detected in the INO80–nucleosome complex. To further improve the reconstruction, we performed 3D flexible refinement in cryoSPARC (46), which yielded an improved map for the A-module and enabled fitting of the AlphaFold3 (47)-predicted A-module structure alongside the observed C-module.

In the hexasome dataset, the A-module binds to unwrapped nucleosomal DNA (**fig. 6a**). In contrast, in the nucleosome dataset, it binds to extranucleosomal DNA (**fig 6b**). Thus, although engaging different DNA sequences, the A-module consistently targets topologically equivalent regions in the H-3 and N-6 states, recognizing extranucleosomal or extrahexasomal DNA, respectively.

Importantly, in the nucleotide-free INO80 structures, we observed clear density for the post-HSA domain (**fig 6b**). This density is consistently absent in structures determined in the presence of ADP-BeF₃, (**fig 6a,d**) suggesting that the post-HSA domain is a dynamic feature in INO80, whose conformation may be coupled to the nucleotide state of the Ino80^motor^, consistent with previous observations in fungal INO80 (31).

Notably, in the H2A.Z nucleosome dataset (with 50 bp flanking DNA on both sides, 50N50), the A-module appears to engage both the entry and exit DNA (**fig 6c,d**). A prior study (30) demonstrated that INO80 can sense both entry and exit DNA to position nucleosomes, a property that likely contributes to its ability to centrally position nucleosomes.

## Discussion

Chromatin remodellers utilize ATP hydrolysis to alter nucleosome structure, composition or position and regulate DNA accessibility. These enzymes are broadly classified into four main families: SWI/SNF, INO80, ISWI, and CHD. While SWI/SNF, ISWI, and CHD remodellers typically bind superhelical location 2 (SHL-2) on the nucleosome, INO80 exhibits a distinct binding mode, as biochemical and structural data have demonstrated from three different species that it binds at SHL-6/-7 on intact nucleosomes—a position notably different from other remodellers (28,29,56–59).

Recent work on yeast and fungal INO80 showed that they can mobilize not only nucleosomes but also hexasomes. The occurrence and role of hexasomes during various DNA associated processes is a developing field (60,61). Hexasomes have been reported to be generated by RNA polymerase II during transcription and can either stall or accelerate elongation depending on whether the promoter-proximal or promoter-distal H2A–H2B dimer is lost (26,62). Moreover, loss of one H2A–H2B dimer creates an asymmetrically accessible substrate for DNA-binding factors and histone-modifying enzymes (63,64). Beyond transcription, hexasomes containing variants such as H2A.Z and H3.3 have been shown to facilitate access of base-excision repair glycosylases to sites of DNA damage, underscoring their versatile, emerging roles in genome maintenance (65).

Given the emerging role of hexasomes as potential substrates but also possible intermediates prompted us to investigate whether human INO80, like its fungal and yeast orthologs, can act on such subnucleosomal species. This is an important question since human INO80 lacks an important part of the ARP5 insertion domain that binds to H2A/H2B acidic patch on the nucleosome, or H3/H4 in the hexasome in case of yeast and fungal INO80 (24,25,29,30). Nevertheless, we find that human INO80 efficiently slides hexasomes. Compared to the nucleosome, the hexasome spin rotates relative to human INO80, resulting in the engagement of Ino80^motor^ at the SHL –2/-3 position. Human INO80 does not, per se, slide hexasomes more efficiently than nucleosomes; when sufficient linker DNA is present for A-module binding, INO80 slides both substrates similarly. This suggests that for human INO80, hexasomes are not per se a preferred substrate over a nucleosome, but may become a good if not preferred substrate if sufficiently long linker DNA is exposed as a result of DNA unwrapping to provide a binding site for the A-module.

Interestingly, we also observed that INO80 can also spin-rotate on the nucleosome, with the Ino80^motor^ bound to either SHL-6 or SHL-7. Comparing both binding modes, the main difference is a partial unwrapping of nucleosomal DNA in the SHL-6 bound state of the ATPase (N-6). Thus, whether bound to nucleosomes or hexasomes, INO80’s ATPase consistently engages DNA at the entry side of the nucleosomal/hexasomal DNA. This binding behavior appears to be remarkably independent of nucleotide presence and nucleosome type. The DNA entry side of the nucleosome is characterized by increased dynamics and accessibility, reduced DNA curvature, and fewer DNA-histone interactions compared to the tightly wrapped core DNA (66–69). This suggests that human INO80 integrates two structural features, DNA curvature around the nucleosome core (ARP5/IES6) and emerging straight DNA at the entry DNA (Ino80^motor^). Since we visualized three spin-rotated binding modes of INO80 from the same species, we can rule out species-specific differences and conclude that the human INO80 (C-module) complex binds nucleosomes in a topology driven manner. The topology driven interaction may equip INO80 with the possibility to act on a range of nucleosomes and also enable regulation through nucleosome and linker DNA features that impact entry DNA paths and dynamics. Remodellers that bind at SHL-2, such as SWI/SNF, ISWI, SWR1and CHD engage DNA where it is already bent by histone interactions, making them less sensitive to dynamic DNA shape features(5,7). This difference highlights INO80’s unique adaptation to exploit nucleosomal DNA flexibility for its specialized roles.

The IES2 subunit is a rather poorly understood, yet conserved component of INO80 remodellers. It’s C-terminal part provides a critical anchor to the RuvBL1/2 while the preceding region helps connect the Ino80^motor^ to the nucleosome gyres via a “throttle” helix (23,24,28,29) contributing to INO80’s ATPase activity and nucleosome mobilization. The functions of the N-terminal region of IES2 have not been revealed. In our study, we made two important observations regarding the function of the N-terminal part of IES2. IES2 contains an RxR motif that binds the acidic patch on the distal side of the nucleosome and this interaction is important for the coupling of ATP hydrolysis steps to the sliding of nucleosomes. IES2 acidic patch binding by IES2 is observed on both nucleosomal binding modes (N-6, N-7), but we did not observe this interaction when bound to hexasomes (H-3), or when bound to the H2A.Z 50N50 nucleosome (NZ-7). The lack of hexasome interactions is consistent with our biochemical observations, since acidic patch mutations affect nucleosome but not hexasome sliding. Human INO80 does not bind the proximal acidic patch via ARP5 grappler, in contrast to fungal INO80 (29,31) therefore IES2 confers the only acidic patch binding of human INO80. In the case of hexasomes, the spin rotation probably sterically prevents IES2 acidic patch binding, while H2A.Z contains three different residue changes compared to canonical H2A within 5Å from IES2 (S9-a), which might explain the reduced binding (**fig S14a**) (37). Thus, proximal acidic patch binding by IES2 could enable INO80 to differentiate between different nucleosomal species, or help retain the proximal H2A/H2B dimer during the sliding reaction in canonical nucleosomes.

Furthermore, we find density for the N-terminal part of IES2 in place of unwrapped nucleosomal DNA at the exit side of the nucleosome. IES2 is the paralog to SWC2 in SWR1 remodellers and shares throttle helix and RuvBL1/2 binding elements (69,70). Like IES2, SWC2 also binds the distal side acidic patch even though SWR1 binds nucleosomes very differently from INO80 (SHL-2) (71). Interestingly, yeast Swc2 has also been shown to partially unwrap DNA from the exit side of the nucleosome (70). It furthermore plays an important role in SWR1’s ability to swap sides on the nucleosome, and has H2A.Z histone chaperone functions (72). Hence, the functional equivalence to SWC2 as histone binding element extends even beyond the acidic patch binding. Our results suggest that IES2 might either help unwrap or stabilize unwrapped exit DNA, with a preference of H2A over H2A.Z. Such a role could prevent H2A/H2B dimer loss during remodelling. Alternatively, IES2 could prepare the exit side of the nucleosome to become the new entry side, if INO80 also has SWR1’s capability to swap sides on a nucleosome (73).

The human INO80 lacks a major part of Arp5 grappler domain, in particular the foot region that is crucial for sliding histones in fungal and yeast INO80, both in the case of nucleosome and hexasome substrates (23,24,29). Instead of a ARP5 grappler interaction, we observed that in the case of human INO80, RuvBL1/2 confer some interactions to the histone cores. Specifically, RuvBL1/2 binds the distal side of histone H2A as the DNA unwraps from the histone specifically in the case of nucleosome and interacts with the phosphate backbone of the unwrapped nucleosomal/hexasomal DNA. In *Ct*INO80, Rvb2 is near the distal side of the H2A/H2B histones when bound to a nucleosome and H3/H4 when bound to a hexasome. However, in *Ct*INO80, we did not observe any interaction of Rvb 1/2 with the histone core as observed in human INO80. Notably, *Ct*INO80’s Rvb-2 E-chain features a positively charged residue (R212) at the corresponding site, leading to electrostatic repulsion rather than interaction (**fig S14b**). These findings suggest that the lack of the grappler domain in human INO80 might be somewhat compensated by the RuvBL1/2, which stabilizes the histone core once the DNA is unwrapped from the nucleosome.

Previous studies have shown that the INO80 ruler mechanism influences the direction of nucleosome sliding, moving it either toward or away from barriers such as DNA ends or neighboring nucleosomes to achieve stable positioning (11,12). In this study, we have shown, that the A-module can interact simultaneously with the entry and exit sides of the nucleosomal DNA. This dual sensitivity is significant as it suggests how A-module may function as a sensor for DNA length. Previous research demonstrated that the chromatin remodeller CHD1 can also sense both the entry and exit sides of DNA (74). This dual sensitivity allows for faster and more precise responses, which is critical for generating regularly spaced nucleosomes. It may function as an autoinhibitory mechanism, slowing nucleosome sliding when extra-nucleosomal DNA is present on the exit side.

In conclusion, our work has illuminated hexasome sliding as a universal capability of INO80. We have also provided high-resolution structures of the INO80 C-module in two distinct states (N-7 and N-6), altogether arguing for a predominantly topological recognition of nucleosomes driven by the degree of entry DNA unwrapping. Exit DNA interactions of the A-module and binding of histones at unwrapped exit DNA by IES2 suggests that INO80’s spectrum of nucleosome recognition and potential remodelling intermediates is far more complex than anticipated from previous work. Studies on more complex nucleosomal substrates are therefore needed to unravel the complex dynamics of INO80 in a chromatin environment.

## Acknowledgments

We thank Katja Lammens and Michael Kugler for help with data collection. We thank Min Zhang for her valuable input in the analysis of remodelling assays, and Mariia Likhodeeva for assistance with nucleosome preparation. We are also grateful to Dirk Kostrewa for his help with model building.

## Data and materials availability

Coordinates for the INO80–hexasome complex (state H-3), the INO80–nucleosome complex (states N-6 and N-7), and the INO80–H2A.Z nucleosome complex (state NZ-7), as well as for the hexasome of the INO80–hexasome complex, the nucleosome of the INO80–nucleosome complex (state N-7 and N-6), and the H2A.Z nucleosome from the INO80–H2A.Z complex (state NZ-7), have been deposited in the Protein Data Bank under accession codes **9GE5, 9GFB, 9GEV, 9GCG, 9GF6,** and **9GE4**. Corresponding cryo-EM density maps have been deposited in the Electron Microscopy Data Bank under accession codes **EMD-51290, EMD-51294, EMD-51310, EMD-51312, EMD-51313, EMD-51229,** and **EMD-51289**. The map for the combined C– and A-modules of the INO80–Nucleosome complex is available in EMDB under accession code **EMD-53574**.

## Funding

This work is supported by the Deutsche Forschungsgemeinschaft DFG (HO 2489/9-1, SFB1064 – 213249687, SFB1361 – 393547839) and the European Research Council (ERC Advanced Grant 833613 INO3D).

## Author contributions

Conceptualization: K.P.H. Construct design: P.A., M.S., and S.W. Cloning: P.A., M.S., S.W., and M.M. Protein purification: P.A., M.S., and F.K. Cryo-EM sample preparation: P.A., and M.S. Data Processing: P.A., M.S., and K.P.H. Model building: P.A., and M.S.*In vitro* biochemistry: P.A., M.S. Visualization: P.A, and M.S. Supervision: K.P.H. Manuscript Writing: K.P.H., P.A., and M.S. with input from the other authors.

## Competing interests

The authors declare no competing interests.

---

**Supplementary figure 1:**
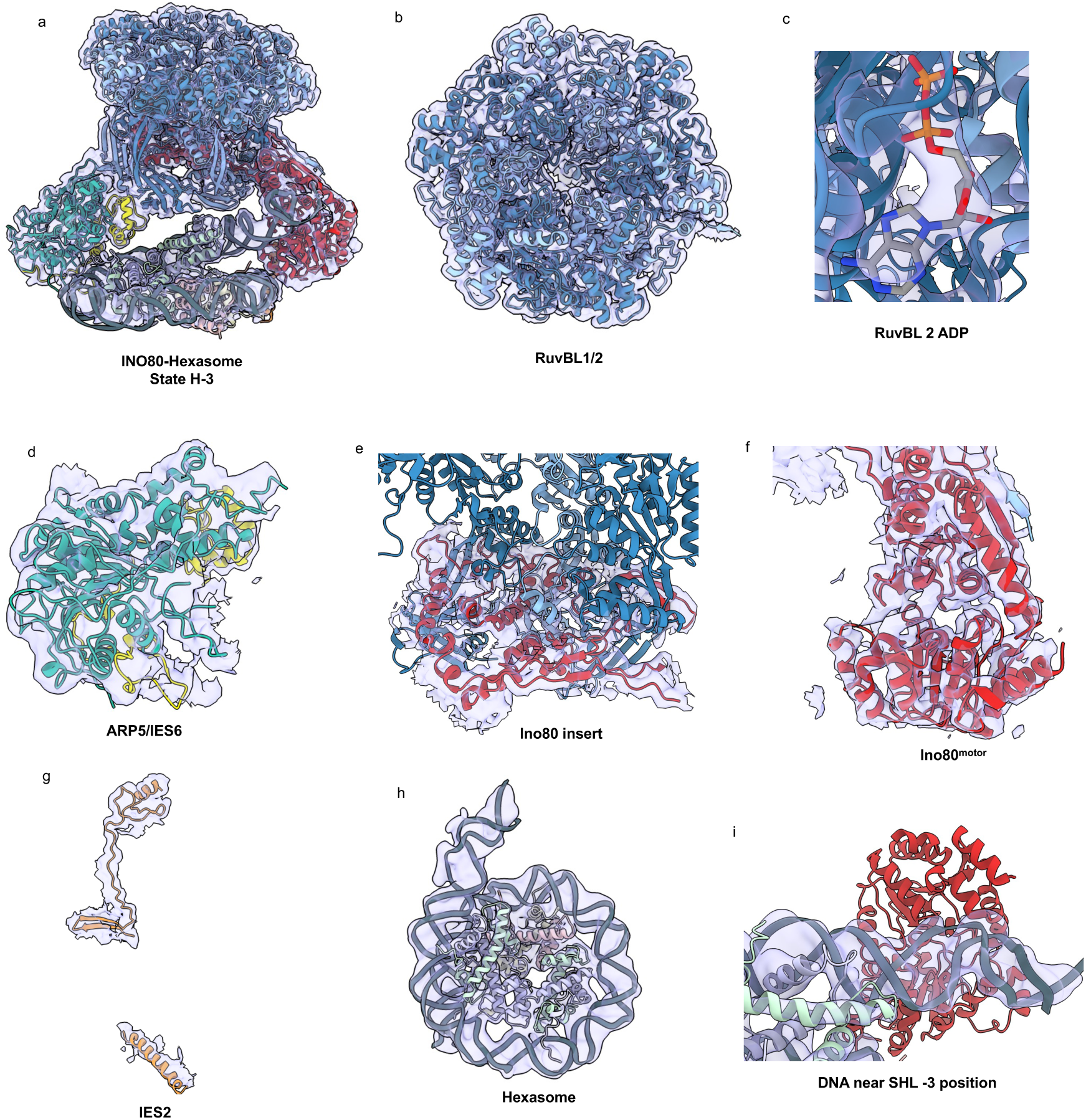
Representative CryoEM density maps and models for INO80-hexasome complex (state H-3) **(a)** Fit of the composite model of the INO80 core module-hexasome complex (state H-3) into the overall density of the whole particle refinement. **(b)** Map density for RuvBL1/2. **(c)** Representative modeled nucleotide (ADP) map densities, observed in all six nucleotide-binding pockets of the RuvBL1/2 heterohexamer. **(d-h)** Representative map densities for ARP5/IES6, the Ino80 insert, Ino80^motor^ and the IES2. **(h)** Map density for the hexasome. **(i)** Representative map density for hexasomal DNA near the SHL-3 position.

**Supplementary figure 2:**
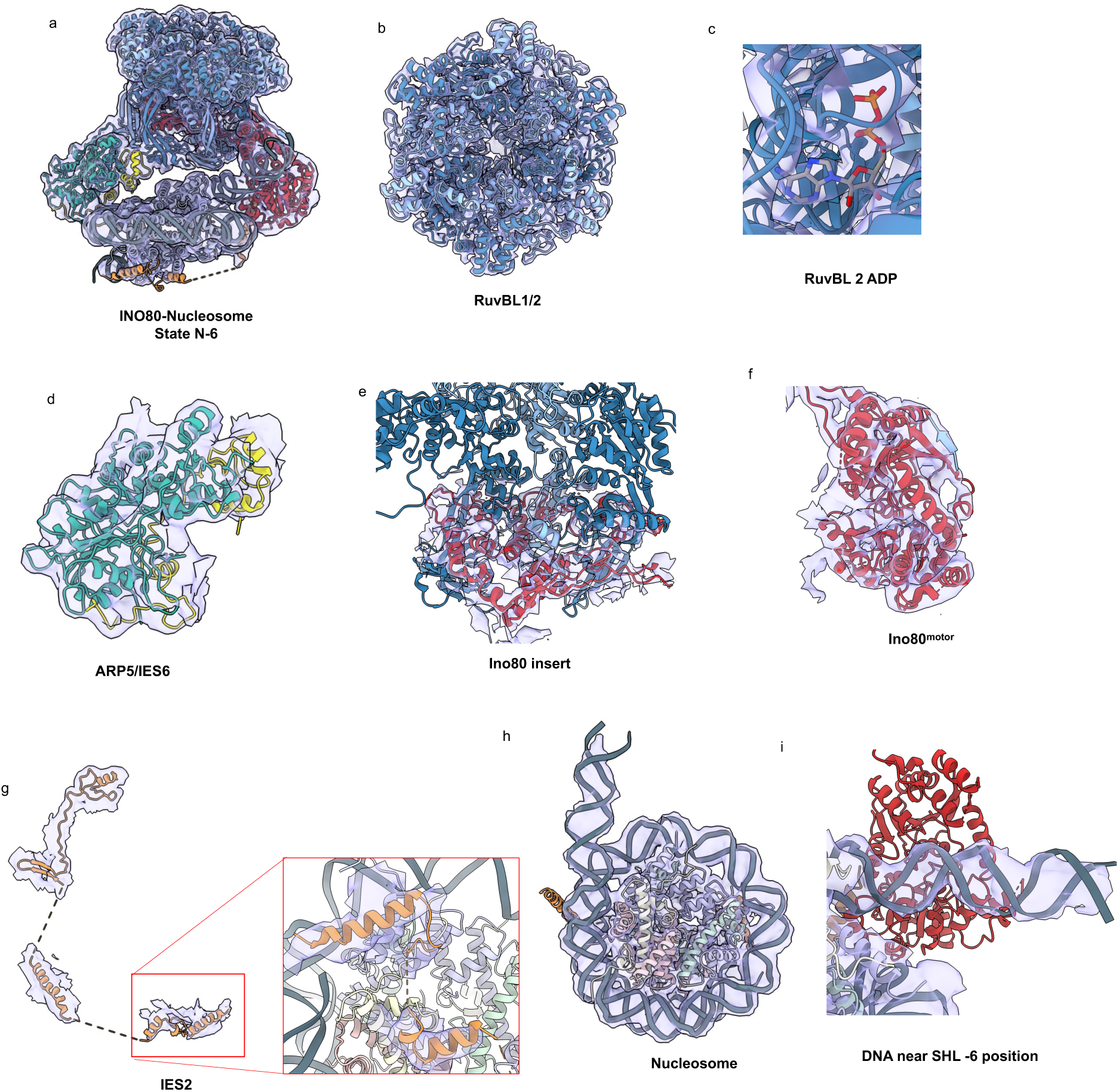
Representative CryoEM density maps and models for INO80-nucleosome complex (state N-6) **(a)** Fit of the composite model of the INO80 C-module-nucleosome complex (state N-6) into the overall density of the whole particle refinement. **(b)** Map density for RuvBL1/2. **(c)** Representative modeled nucleotide (ADP) map densities, observed in all six nucleotide-binding pockets of the RuvBL1/2 heterohexamer. **(d-f)** Representative map densities for ARP5/IES6, the Ino80 insert, and the Ino80^motor^. **(g)** Overall map density for IES2, with a focused representation of the IES2 region (G140-S189), which interacts with the distal side of the nucleosome. **(h, i)** Map density for the nucleosome (state N-6). **(i)** Representative map density for nucleosomal DNA near the SHL-6 position.

**Supplementary figure 3:**
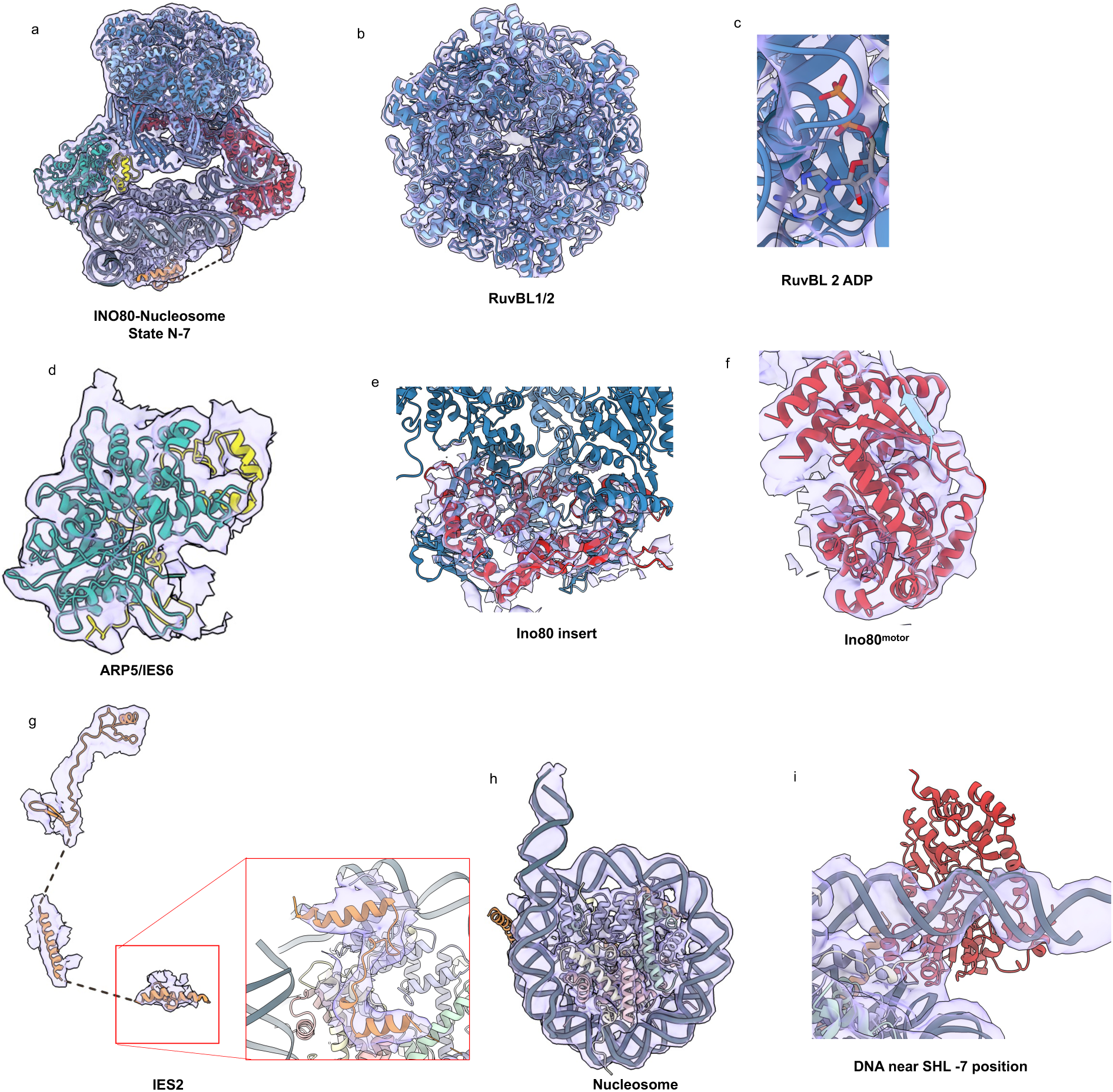
Representative CryoEM density maps and models for INO80-nucleosome complex (state N-7) **(a)** Fit of the composite model of the INO80 C-module-nucleosome complex (state N-7) into the overall density of the whole particle refinement. **(b)** Map density for RuvBL1/2. **(c)** Representative modeled nucleotide (ADP) map densities, observed in all six nucleotide-binding pockets of the RuvBL1/2 heterohexamer. **(d-f)** Representative map densities for ARO5/IES6, the Ino80 insert, and the Ino80^motor^. **(g)** Overall map density for IES2, with a focused representation of the IES2 region (G140-S189), which interacts with the distal side of the nucleosome **(h, i)** Map density for the nucleosome (state N-7). **(i)** Representative map density for nucleosomal DNA near the SHL-6 position.

**Supplementary figure 4:**
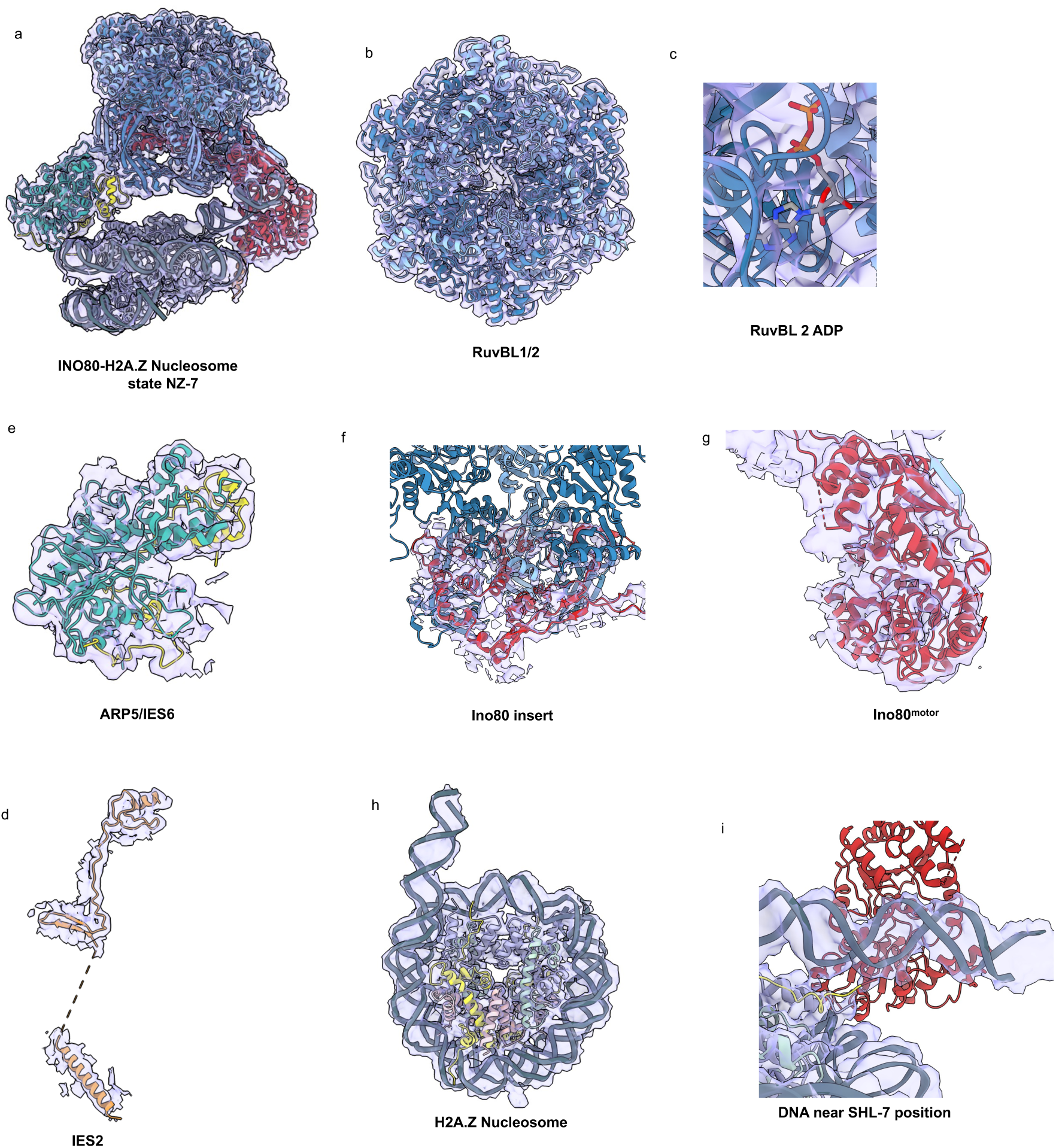
Representative CryoEM density maps and models for INO80-H2A.Z nucleosome complex (state NZ-7) (**a**) Fit of the composite model of the INO80 C-module-H2A.Znucleosome complex (state NZ-7) into the overall density of the whole particle refinement. (**b**) Map density for RuvBL1/2. (**c**) Representative modeled nucleotide (ADP) map densities, observed in all six nucleotide-binding pockets of the RuvBL1/2 heterohexamer. (**d-g**) Representative map densities for ARP5/IES6, the Ino80 insert, Ino80^motor^ and the IES2, (**h**) Map density for the H2A.Z nucleosome. (**i)** Representative map density for nucleosomal DNA near the SHL-7 position.

**Supplementary figure 5:**
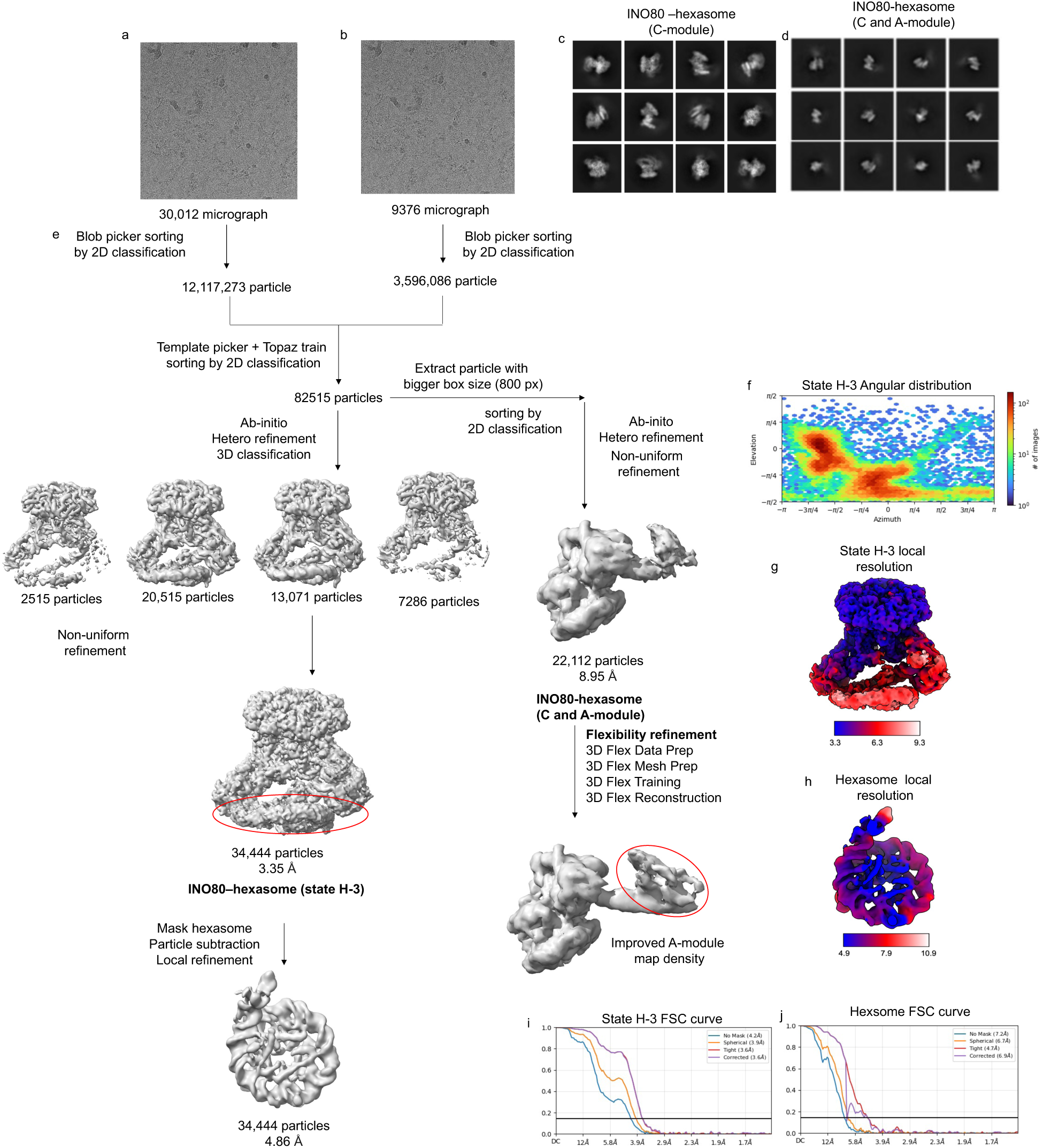
Cryo-EM refinement procedures of the INO80-hexasome complex. (**a,b**) Representative micrographs of the INO80–hexasome complex from two different data collections. (**c**) Representative classes from the 2D classification of particles used for the final INO80–hexasome reconstruction. (**d**) Representative classes from the 2D classification of particles used for the final reconstruction of the INO80-hexasome (C and A-module). (**e**) Workflow schematic illustrating the steps used to obtain high-quality maps of the core–hexasome and the INO80–hexasome (C– and A-module). (**f**) Angular distribution plot of particles employed to reconstruct map for the state H-3. (**g**) Local resolution map of the state H-3. (**h**) Local resolution map of the hexasome. (**i**) FSC curves of the state H-3 map. Resolution at the FSC threshold criterion 0.143 is indicated. (**j)** FSC curves of the hexasome. Resolution at the FSC threshold criterion 0.143 is indicated.

**Supplementary figure 6:**
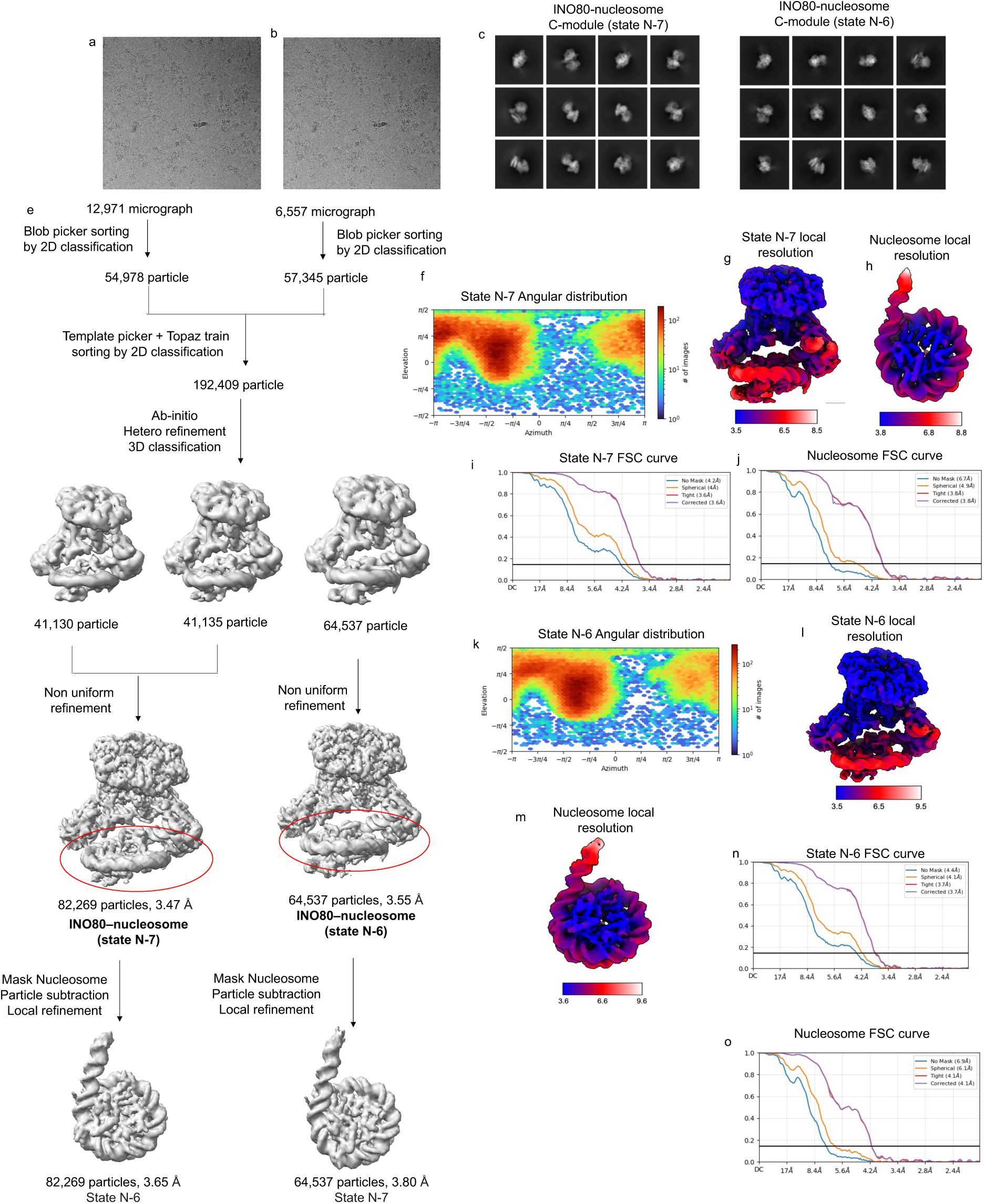
Cryo-EM refinement procedures of the INO80-nucleosome complex. (**a, b**) Representative micrographs of the INO80–nucleosome complex from two different data collections. (**c**) Representative classes from the 2D classification of particles used for the final INO80–nucleosome reconstruction for state N-7. (**d**) Representative classes from the 2D classification of particles used for the final INO80–nucleosome reconstruction for state N-6. (**e**) Workflow schematic illustrating the steps to obtain a high-quality core–hexasome map. (**f**) Angular distribution plot of particles employed to reconstruct map for the state N-7. (**g**) Local resolution map of the state H-3 (**j**)Local resolution map of the nucleosome (state N-7). (g) FSC curves of the state N-7 map. Resolution at the FSC threshold criterion 0.143 is indicated (i) FSC curves of the nucleosome. Resolution at the FSC threshold criterion 0.143 is indicated (**k**) Angular distribution plot of particles employed to reconstruct map for the state N-6. (**l**) Local resolution map of the state N-6 (**m)** Local resolution map of the nucleosome (state N-6) (**n**) FSC curves of the state N-7 map. Resolution at the FSC threshold criterion 0.143 is indicated. (**o**) FSC curves of the nucleosome (state N-6). Resolution at the FSC threshold criterion 0.143 is indicated.

**Supplementary figure 7:**
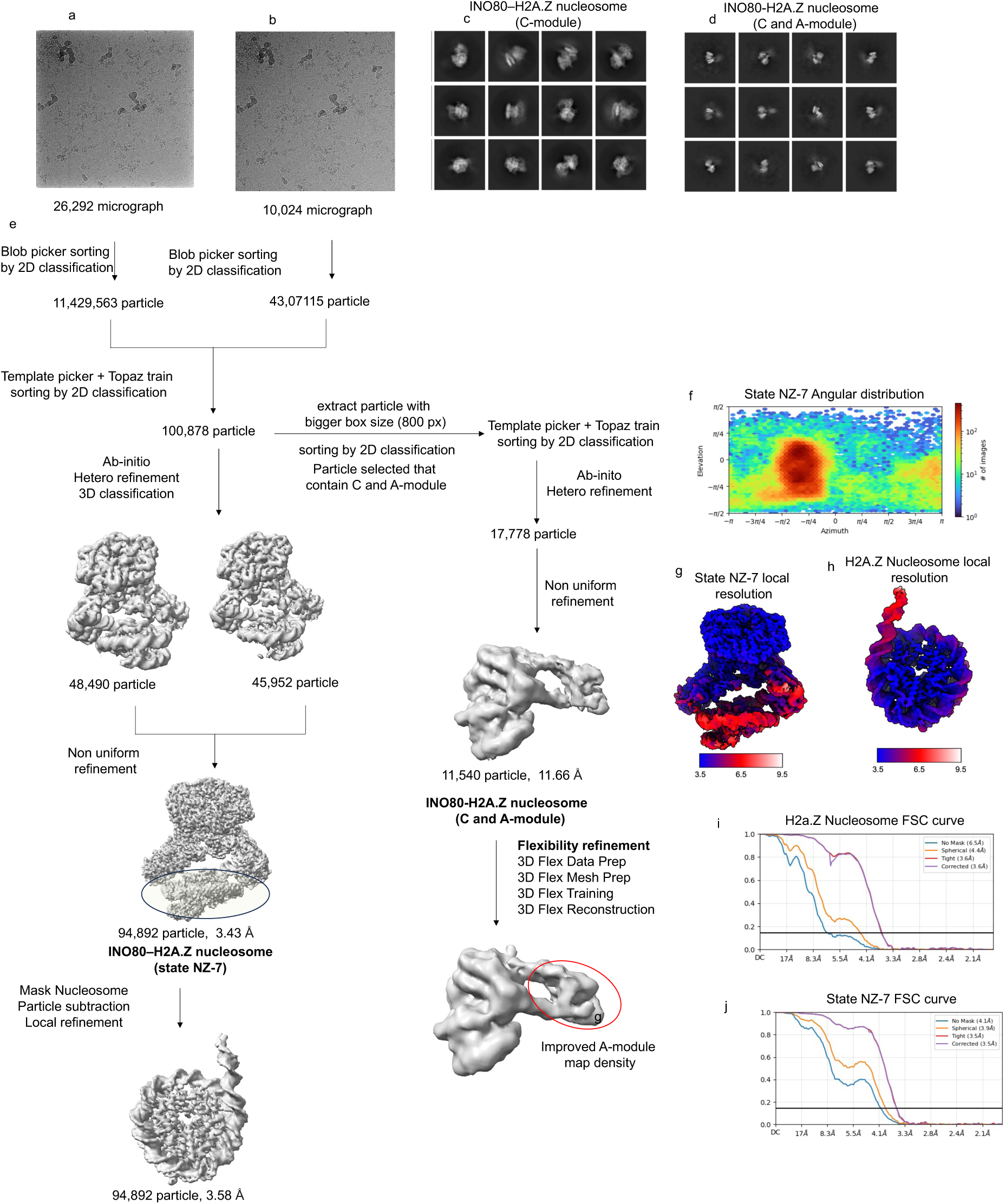
Cryo-EM refinement procedures of the INO80-H2A.Z nucleosome complex. (**a, b**) Representative micrographs of the INO80–H2A.Z nucleosome complex from two different data collections.(**c**) Representative classes from the 2D classification of particles used for the final INO80–H2A.Z nucleosome (state NZ-7) reconstruction.(**d**) Representative classes from the 2D classification of particles used for the final INO80-H2A.Z nucleosomse (C and A module) reconstruction.(**e**) Workflow schematic illustrating the steps to obtain a high-quality core–hexasome map.(**f**) Angular distribution plot of particles employed to reconstruct map for the state NZ-7. indicated. **(g)** Local resolution map of the state NZ-7. **(h)** Local resolution map of the H2A.Z nucleosome. **(i)** FSC curves of the state NZ-7 map. Resolution at the FSC threshold criterion 0.143 is indicated. **(j)** FSC curves of the H2A.Z nucleosome. Resolution at the FSC threshold criterion 0.143 is indicated.

**Supplementary figure 8:**
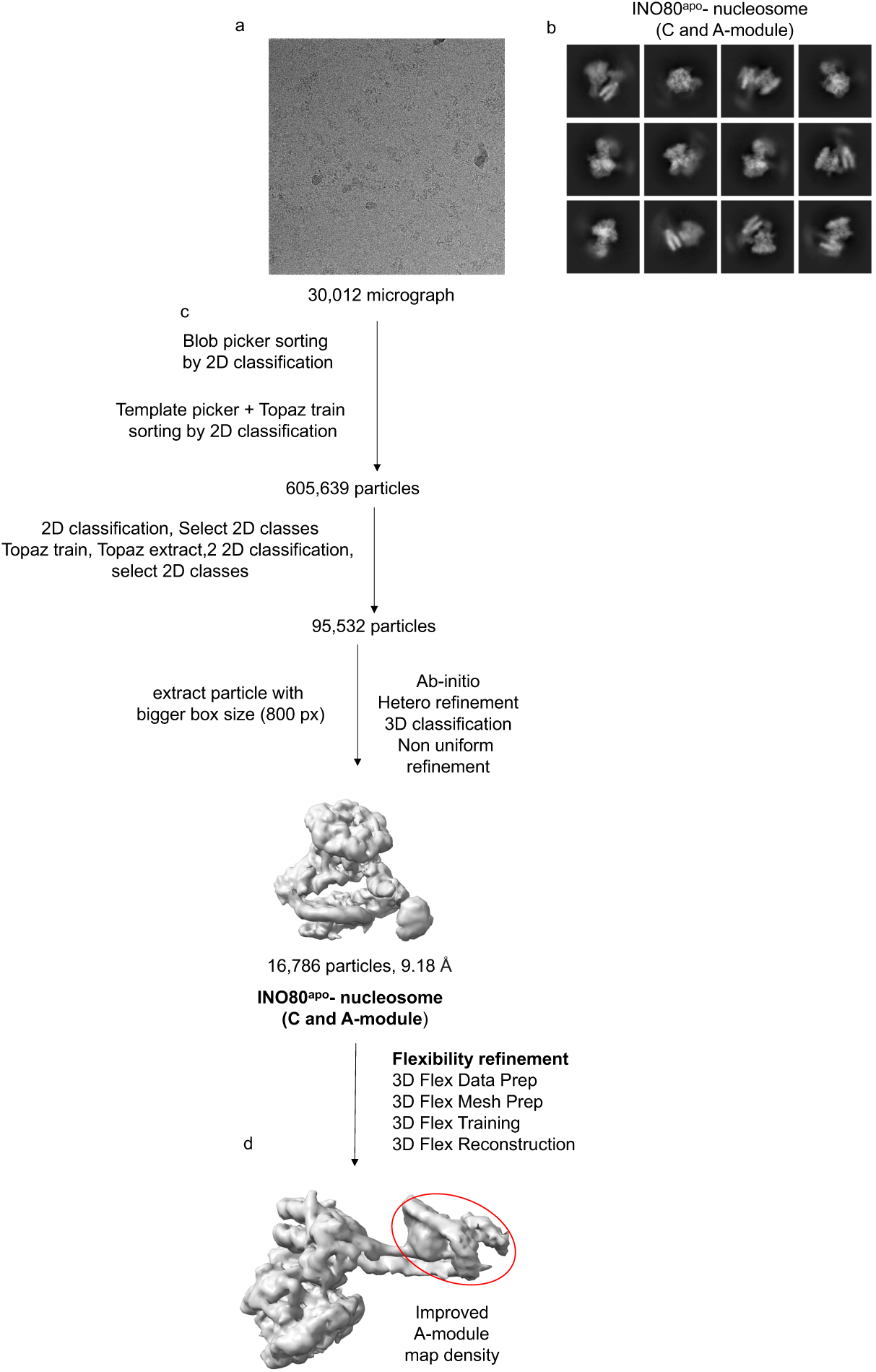
Cryo-EM refinement procedures of the INO80^apo^-Nucleosome (C and A-module) complex **(a)** Representative micrographs of the INO80^apo^ –nucleosome (C and A module) complex. **(b)** Representative classes from the 2D classification of particles used for the final INO80^apo^–nucleosome C and A-module complex reconstruction. **(c)** Workflow schematic illustrating the steps to obtain INO80^apo^-nucleosome (C and A module) map. **(d)** 3D flexible refinement (3D-flex) was performed to enhance map quality for the flexible A-module.

**Supplementary figure 9:**
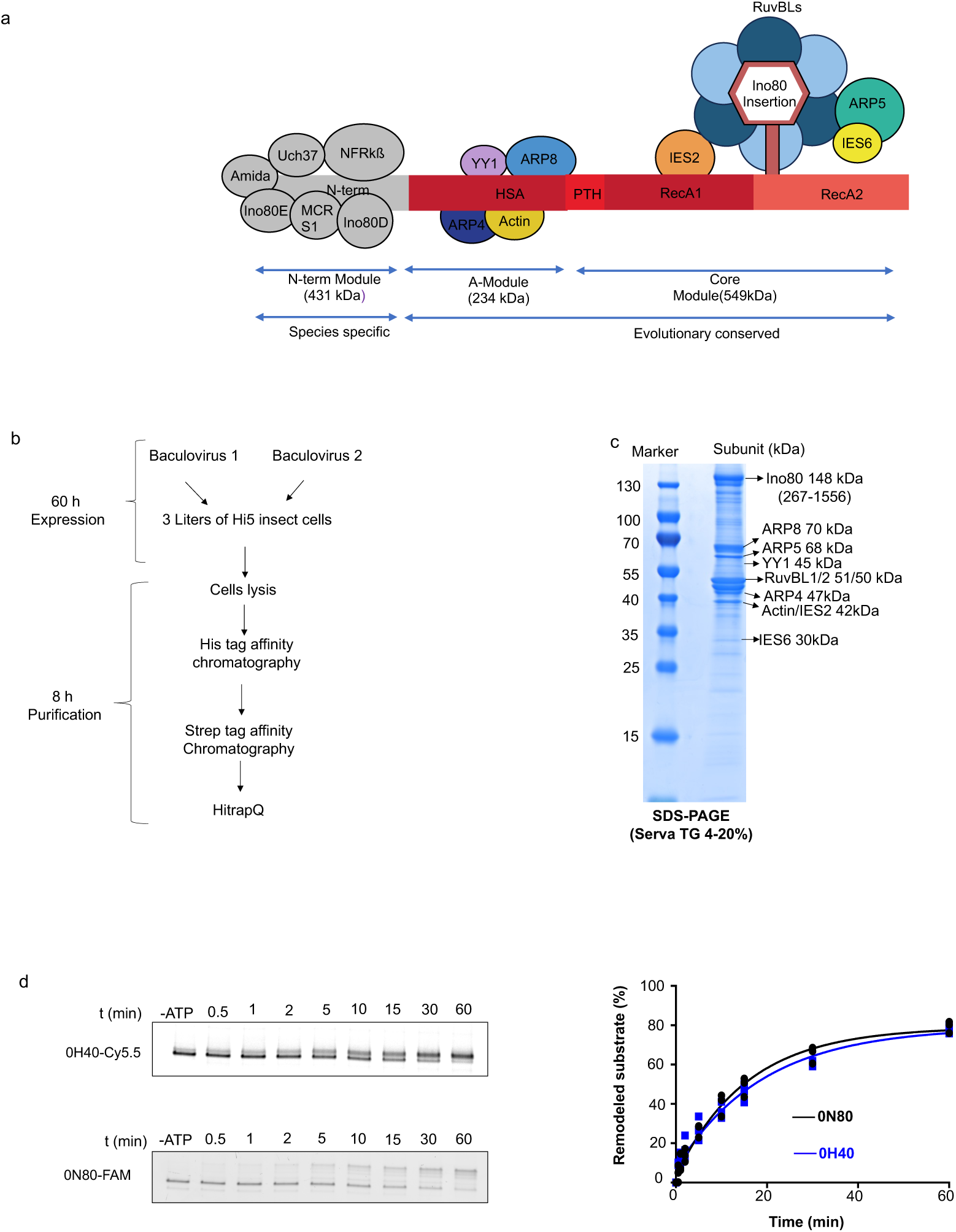
(a) Schematic of the subunit composition of INO80. The N-terminally truncated (delta-N) INO80 is used in this study. Subunits are categorized into the INO80 C-module and A-module. **(b)** Protein expression and purification strategy used for wild type (delta-N) and mutant INO80. **(c)** SDS-PAGE of the purified wild type delta-N INO80 (staining was done with Coomassie Brilliant Blue R250). The identity of each protein complex band was confirmed by Mass spectrometry (supplementary excel file). **(d**) The native PAGE analysis and evaluation of the competition sliding activity of INO80 on nucleosomes and hexasomes. The activities of INO80 on both nucleosome and hexasome substrates were monitored in the same reaction tube. Nucleosome 0N80 (601 DNA + 80 bp) and hexasome 0H40 (601 DNA + 40 bp) were selected to maintain the same linker length, as ∼40 bp of DNA becomes unwrapped upon the formation of hexasome. The nucleosome 0N80 and hexasome 0H40 were labeled with the fluorophores FAM and Cy5.5, respectively. The native PAGE was scanned with both fluorophores’ signals. INO80 showed a slight preference for nucleosome over hexasome, keeping the linker length same at 80 bp.

**Supplementary figure 10:**
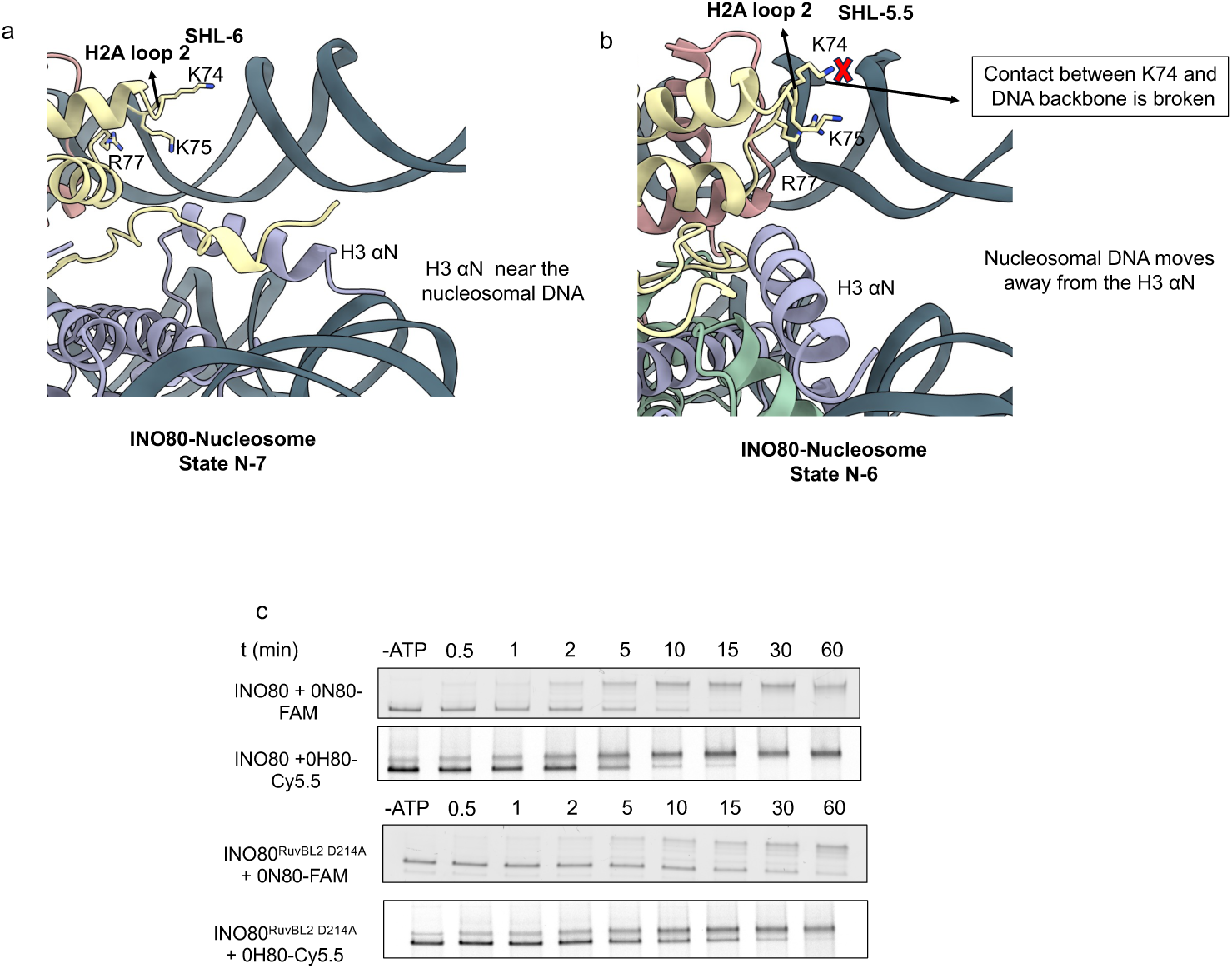
(**a, b**) In state N-7, the proximal-side histone H3 αN and H2A loop 2 are in close contact with nucleosomal DNA near the SHL-6 position. Upon movement of the Ino80^motor^ in state N-6, these contacts are lost, leading to partial unwrapping of DNA and partial exposure of the H2A/H2B dimer. (**c**) Native PAGE analysis of nucleosome sliding assays using end-positioned nucleosomes (6-FAM-labeled) and hexasomes (Cy5.5-labeled) with wild-type INO80 and the INO80 RuvBL2 D214A mutant.

**Supplementary figure 11:**
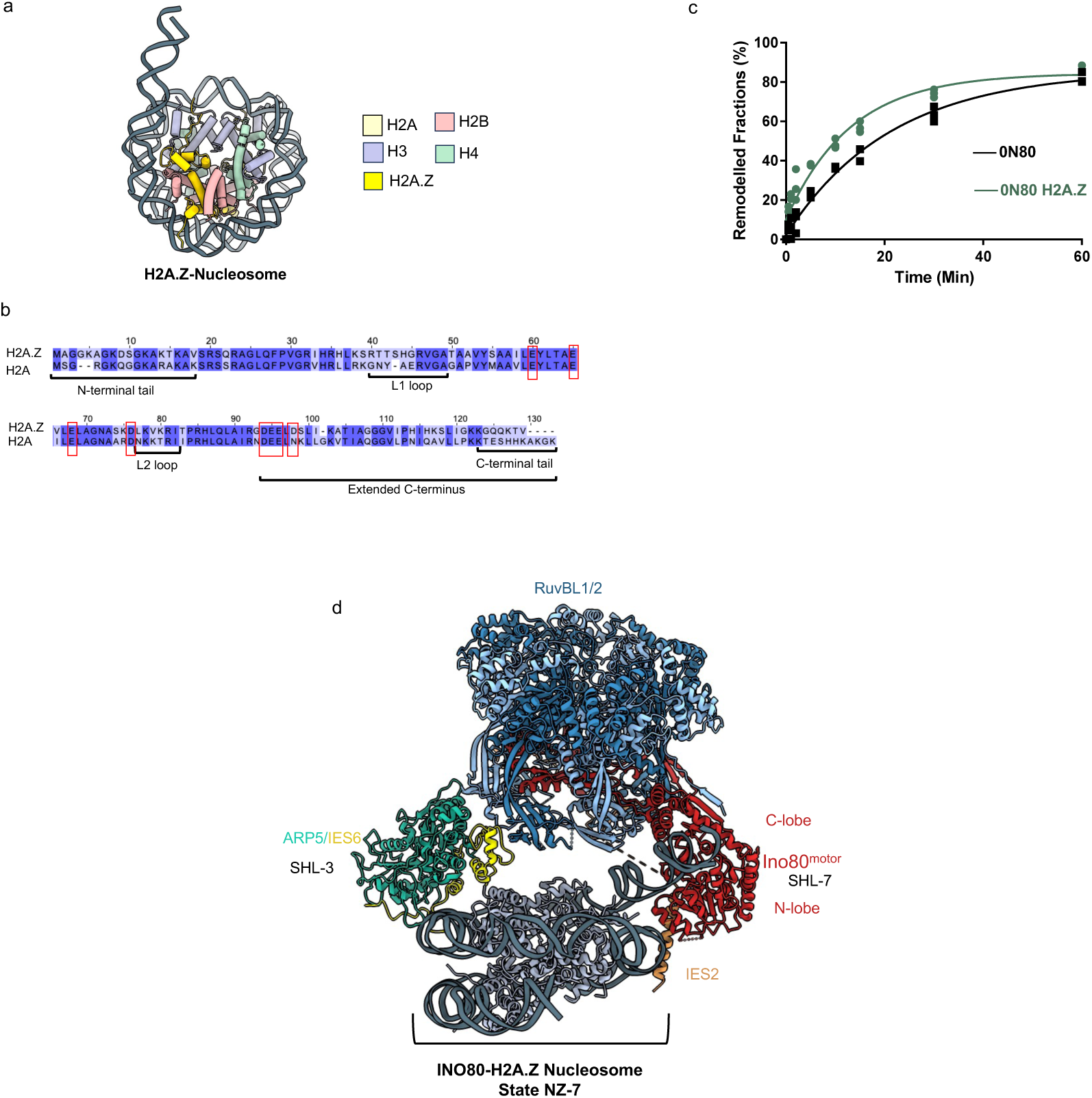
(**a**) Cartoon depiction of the H2A.Z-containing nucleosome from this study. (**b)** Sequence alignment of *H.s., Homo sapiens* H2A and H2A.Z histones, highlighting the acidic patch residues (indicated by a red box). The H2A.Z histone contains an additional acidic residue in its acidic patch region compared to H2A. (**c**) Native PAGE analysis and competitive sliding assay of INO80 activity on H2A.Z nucleosomes and canonical H2A nucleosomes. The H2A.Z nucleosome (0N80) and canonical nucleosome (0N80) were labeled with the fluorophores FAM and Cy5.5, respectively. Native PAGE analysis detected signals from both fluorophores, revealing that INO80 preferentially remodels H2A.Z nucleosomes over canonical nucleosomes. (**d**) Cryo-EM structure of the INO80 C-module bound to the centrally positioned 50N50 H2A.Z variant-containing nucleosome (State NZ-7). The ATPase motor domain interacts with the H2A.Z nucleosome near the SHL-7 position, while on the opposite side, the ARP5/IES6 subunits interact near the SHL-3 position.

**Supplementary figure 12:**
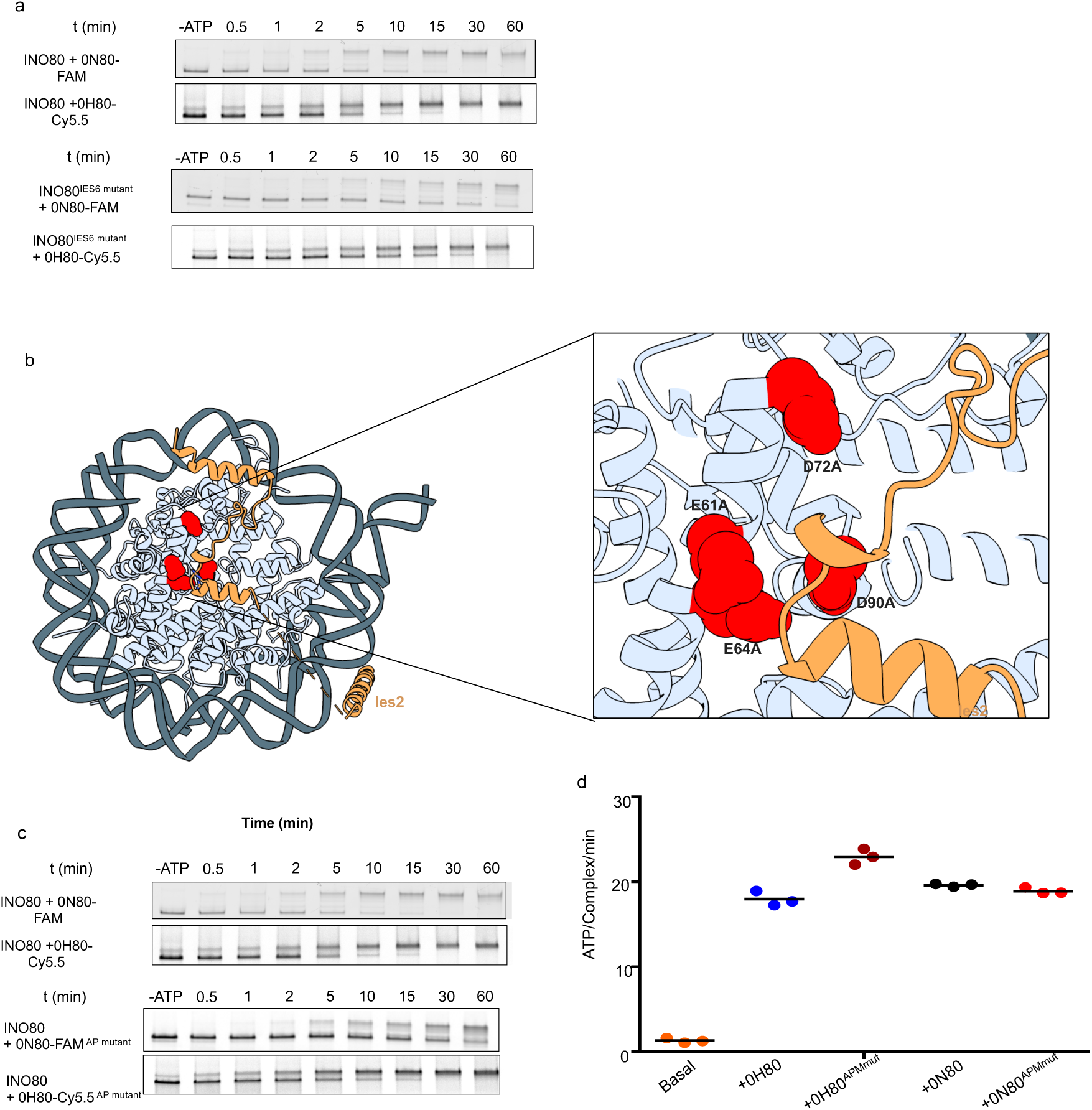
(**a**) Native PAGE analysis of nucleosome sliding assays using end-positioned nucleosomes (FAM-labeled) and hexasomes (Cy5.5-labeled) by wild-type INO80 and its mutant INO80 IES6 (135–137) GKK-to-AAA mutant **(b)** A close-up view of the acidic patch residues (shown in red) on the distal side of the nucleosome, which are mutated on histone H2A.**(c)** Translocation assays of INO80 on hexasomes and nucleosomes containing mutated histone H2A**(d)** Native PAGE analysis showing sliding of end-positioned nucleosomes (FAM-labeled) and hexasomes (Cy5.5-labeled) by INO80. Wild-type nucleosomes (black) and hexasomes (blue) are compared to their respective acidic patch mutants: nucleosome acidic patch mutant (red) and hexasome acidic patch mutant (brick red) **d**) ATPase rates of INO80 were measured with stimulation by mutated hexasome (0H80APmut) and nucleosome (0N80APmut). Control reactions included: basal ATPase activity (no nucleosome/hexasome), wild-type hexasomes (0H80), and wild-type nucleosomes (0N80). ATPase rates were calculated from the linear region of the raw data and corrected using a buffer blank. Mean values and individual data points (n = 3, technical replicates) are shown.

**Supplementary figure 13:**
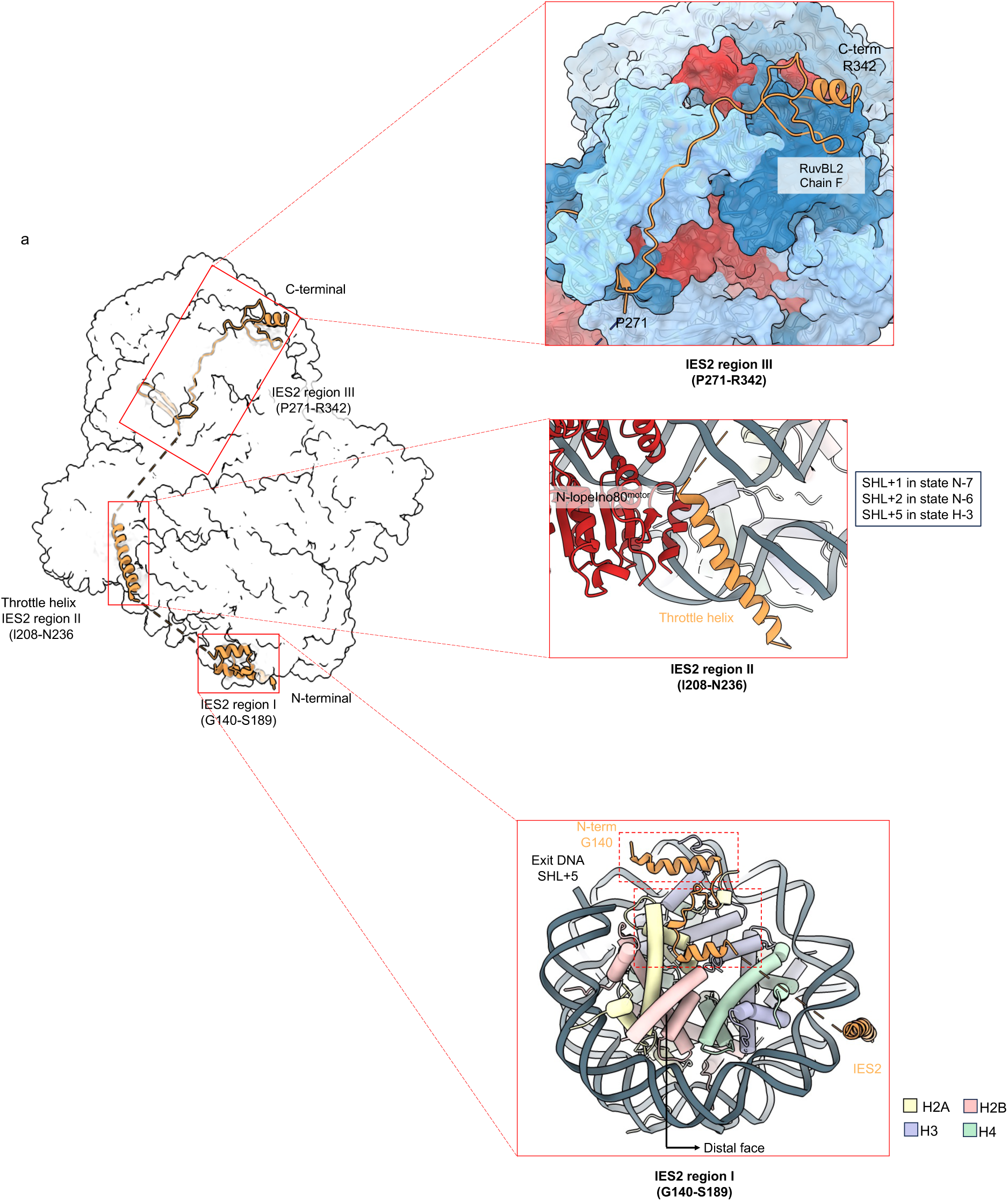
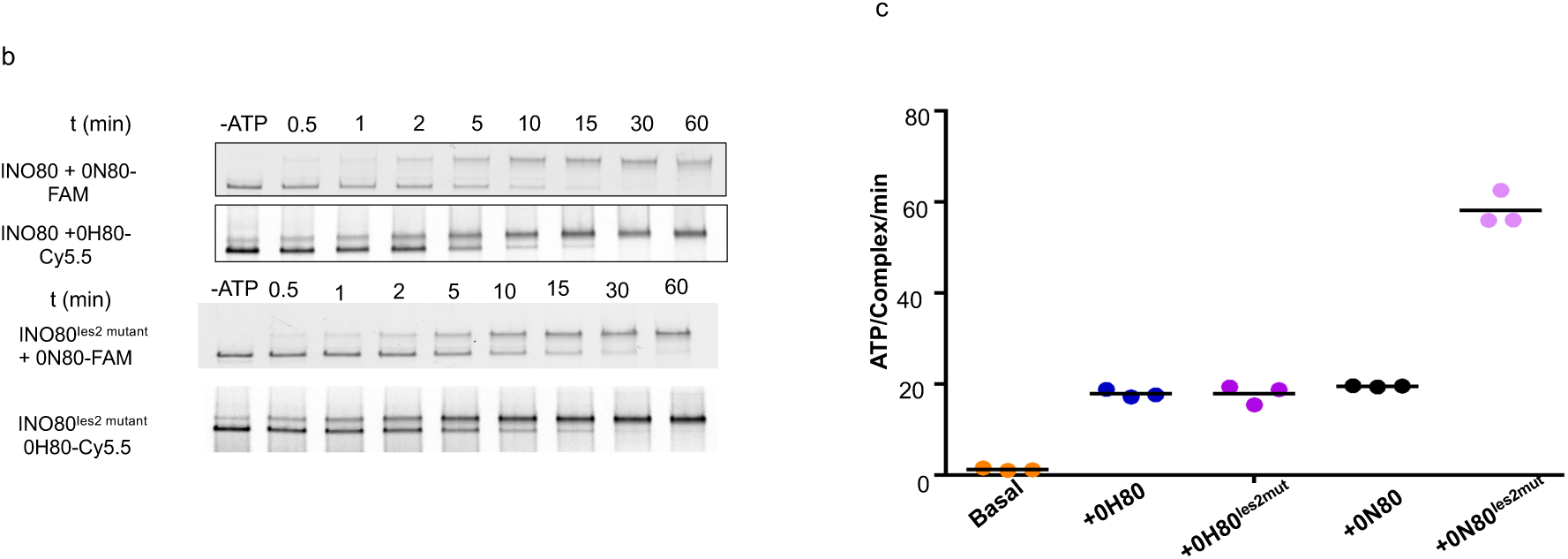
(**a**) The structural model of IES2 (orange), divided into three regions (I, II, III), is shown alongside the hINO80-nucleosome complex (white). Close up view of the IES2 interacts with the acidic patch residues on the distal side of the nucleosome, the ATPase motor domain via its throttle helix, and RuvBL1/2. Interaction of residues G140-S189 (Region I) of IES2 with the histone core of the nucleosome (shown is the State N-7). These interactions are also present in state N-6 (not shown). Close-up view of the IES2 throttle helix (Region II) interacting with the Ino80^motor^ and nucleosomal DNA at the major groove at different SHL positions: SHL+1 in state N-7, SHL+2 in state N-6, and SHL+5 in state H-3. Close-up view of the IES2 Region III interacting with RuvBL1/2. (**b**) Native PAGE analysis of nucleosome sliding assays using end-positioned nucleosomes (FAM-labeled) and hexasomes (Cy5.5-labeled) by wild-type *Hs*INO80 and its mutant INO80 Ies2 mutant. **(c)** ATPase activity of INO80 IES2 mutants, with and without stimulation by nucleosomes or hexasomes. ATPase rates were calculated from the linear region of the raw data and corrected using a buffer blank. Mean values and individual data points (n = 3, technical replicates) are presented.

**Supplementary figure 14:**
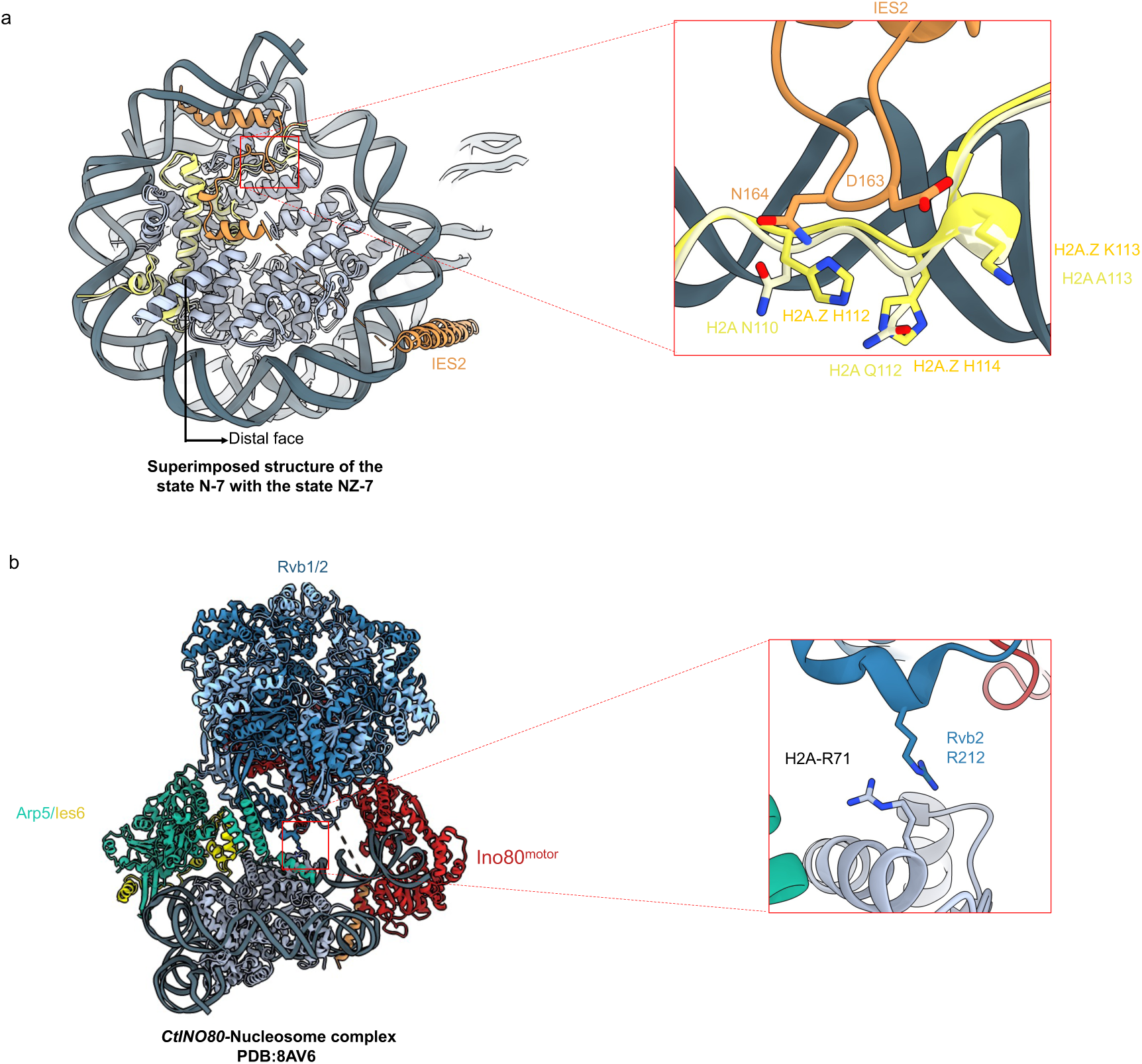
(**a**) Superimposed structure of the INO80-canonical nucleosome state N-7 with the INO80-H2A.Z nucleosome state NZ-7 structure, showing IES2 hairpin loop region residues (D163, N164) interacting with specific H2A.Z residues (K113, H114, H114), along with the corresponding residues present in canonical H2A. **(b)** In the *Ct*INO80-nucleosome complex, similar to the *Hs*INO80 structure, RuvBL2 (referred to as Rvb1/2 in *Ct*INO80) is also positioned near the histone core. However, instead of the D residue present in *Hs*INO80, R212 is found near the histone core in *Ct*INO80. These positively charged residues repel each other, preventing the formation of any interactions.

## Notes

### Competing Interest Statement

The authors have declared no competing interest.

### Summary of Updates

Revised manuscript with updated main figures and revised Supplementary Figure 12 and order of Supplementary figures

